# Regulation of alternative polyadenylation isoforms of *Timp2* is an effector event of RAS signaling in cell transformation

**DOI:** 10.1101/2024.09.26.613909

**Authors:** Yuxi Ai, Qingbao Ding, Zezhong Wan, Sanjay Tyagi, Alexandra Indeglia, Maureen Murphy, Bin Tian

**Affiliations:** Biochemistry and Molecular Biophysics Graduate Group, University of Pennsylvania Perelman School of Medicine, Philadelphia, PA 19104, USA; Department of Biochemistry and Biophysics, University of Pennsylvania Perelman School of Medicine, Philadelphia, PA 19104, USA; Genome Regulation and Cell Signaling Program, The Wistar Institute, Philadelphia, PA 19104, USA; Public Health Research Institute and Department of Medicine, New Jersey Medical School, Rutgers University, Newark, NJ 07103, USA; Molecular and Cellular Oncogenesis Program, The Wistar Institute, Philadelphia, PA 19104, USA

**Keywords:** Timp2, Alternative polyadenylation, Ras, 3’UTR, protein secretion

## Abstract

Alternative polyadenylation (APA) generates mRNA isoforms with different lengths of the 3’ untranslated region (3’ UTR). The tissue inhibitor of metalloproteinase 2 (TIMP2) plays a key role in extracellular matrix remodeling under various developmental and disease conditions. Both human and mouse genes encoding TIMP2 contain two highly conserved 3’UTR APA sites, leading to mRNA isoforms that differ substantially in 3’UTR size. APA of *Timp2* is one of the most significantly regulated events in multiple cell differentiation lineages. Here we show that *Timp2* APA is highly regulated in transformation of NIH3T3 cells by the oncogene HRAS^G12V^. Perturbations of isoform expression with long 3’UTR isoform-specific knockdown or genomic removal of the alternative UTR (aUTR) region indicate that the long 3’UTR isoform contributes to the secreted Timp2 protein much more than the short 3’UTR isoform. The short and long 3’UTR isoforms differ in subcellular localization to endoplasmic reticulum (ER). Strikingly, *Timp2* aUTR enhances secreted protein expression but no effect on intracellular proteins in reporter assays. Furthermore, downregulation of *Timp2* long isoform mitigates gene expression changes elicited by HRAS^G12V^. Together, our data indicate that regulation of Timp2 protein expression through APA isoform changes is an integral part of RAS-mediated cell transformation and 3’UTR isoforms of *Timp2* can have distinct impacts on expression of secreted vs. intracellular proteins.

## INTRODUCTION

Almost all protein-coding transcripts in eukaryotic cells contain a poly(A) tail at its 3’ end, which results from the cleavage and polyadenylation (CPA) reaction acting on the precursor RNA. The CPA site, also known as the polyadenylation site (PAS), is defined by a set of adjacent RNA motifs, the best known of which is the upstream A[A/U]UAAA hexamer or a close variant ^1^. More than 70% of protein-coding genes in the human genome have been found to harbor multiple PASs ^2,3^,leading to expression of mRNA isoforms with different 3’ ends ^4^, a phenomenon known as alternative polyadenylation (APA). Most APA sites are located in the 3’ untraslated regions (3’UTR) in the last exon of a gene, which results in mRNA isoforms with different sizes of 3’UTRs. Accordingly, APA sites proximial to the 5’ end of 3’UTR, or pPAS, lead to short 3’UTR isoforms whereas distal APA sites, or dPAS, lead to long 3’UTR isoforms.

APA isoform expression often exhibits a strong cell type specificity and substantial variation during cell differentiation and development ^5^. For example, widespread 3’UTR shortening events have been reported in various cancer types ^6^ and in secretory cell differentiation ^7^. By contrast, 3’UTRs globally lengthen via APA during mouse embryonic development ^8^ and many cell differentiation lineages, such as myogenesis ^9^ and neurogenesis ^10^. Due to the difference in 3’UTR sequence, APA isoforms are generally believed to have distinct mRNA metabolism, such as mRNA stability, translation, and subcellular localization ^4^.

RAS proteins are GTPases that can bind to GDP in its inactive form but GTP in its active form ^11^. They play important roles in cell growth, proliferation, and migration ^12^. Mutations of RAS genes are common drivers of human cancer ^13^. Mutations in the three canonical RAS genes, namely, *KRAS*, *HRAS* and *NRAS*, have been associated with distinct sets of cancers. For example, *KRAS* mutations have been found in more than 90% of pancreatic cancer patients ^14^; *NRAS* is mutated in 28% of cutaneous melanomas ^15^; *HRAS* is frequently mutated in bladder ^16^ and thyroid ^17^ cancers. Common oncogenic mutations of *RAS* genes are gain-of-function mutations that involve a single amino acid change in the RAS protein that increases the binding affinity of RAS protein to GTP and hence keeps the protein in its active state. As a result, downstream signaling events are constitutively activated. Oncogenic phenotypes associated with *RAS* mutations include uncontrolled cell growth and enhanced cell motility ^18^. The mouse fibroblast cell line NIH3T3 has been an key model to study RAS-mediated cell transformation for many decades ^19^.

Whereas cancer cells have been shown to display global APA-mediated 3’UTR shortening, leading to up-regulation of mRNA expression of many proto-oncogenes ^6,20–22^ and 3’UTR size change can be a molecular biomarker with prognostic potentials for cancers ^23^, little is known about how RAS signaling changes 3’UTR size via APA. One study found that a gain-of-function mutation of let-60 in C. elegans, which encodes the homolog of human RAS protein, causes a multivulva phenotype ^24^. Importantly, depletion of the CPA machinery factor *cfim-1*, the homolog of human *NUDT21* gene, increases the multivulva phenotype. Because *NUDT21* encodes the CFI-25 protein, a major regulator of 3’UTR APA ^9,25^, it is conceivable that RAS signaling and 3’UTR regulation have some functional interactions. In the current study, we examine APA regulation in RAS-transformed NIH3T3 cells. We further analyze the consequences of APA of one key event, i.e., regulation of *Timp2* gene, which encodes tissue inhibitor of metalloproteinase 2 (Timp2), a protein with important functions in cell migration and extracellular matrix remodeling.

## RESULTS

### Transformation of NIH3T3 cells by HRAS^G12V^ leads to downregulation of *Timp2* long isoform and protein expression

To understand alternative polyadenylation (APA) regulation by cell transformation, we overexpressed a gain-of-function mutant form of HRAS (HRAS^G12V^) ^26–29^ in mouse fibroblast NIH3T3 cells, a well-established cell transformation system used in many studies ^30,31^. We constructed a lentiviral vector that expresses HRAS^G12V^ **(Figs. 1A and S1A)** and transduced NIH3T3 cells with the virus. Compared to the cells transduced with control viruses (empty lentiviral vector), the cells overexpressing HRAS^G12V^ had a lower proliferation rate **(Fig. S1B)**, readily formed cell foci in culture dishes **(Fig. S1C)**, and obtained a greater migration ability in wound healing assays **(Fig. S1D)**. These phenotypes are in line with oncogenic RAS-transformed NIH3T3 cells reported before^32,33^, confirming the success of our cell transformation approach.

**Figure 1.**
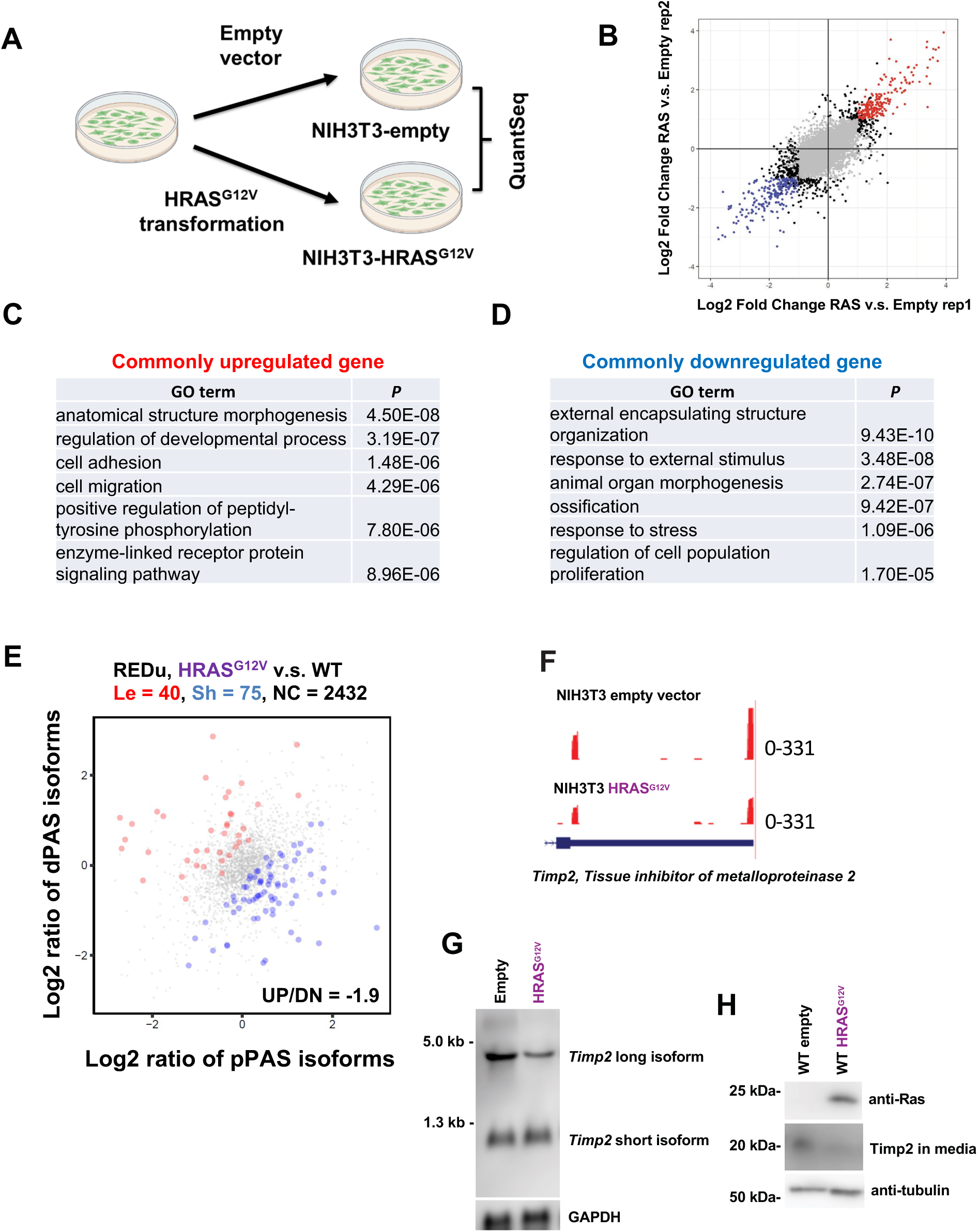
*Timp2* is significantly regulated in gene expression and APA in HRAS^G12V^-transformed NIH3T3 cells. **(A)** NIH3T3 cells were transduced with a lentiviral vector containing an HRAS^G12V^ coding region (NIH3T3-HRAS^G12V^). An empty (NIH3T3-empty) vector was used as a control. Stable cell lines were subject to RNA sequencing by using the QuantSeq-Pool method. **(B)** Differential gene expression analysis of NIH3T3-HRAS^G12V^ vs. NIH3T3-empty cells. A scatter plot shows the correlation of gene expression changes between two biological replicates. Colored genes are those showing P < 0.05 and log2 fold change > 1 in both replicates, with red for commonly upregulated and blue for commonly downregulated. **(C-D)** Gene ontology (GO) analysis result for the commonly upregulated genes (C) and commonly downregulated genes (D), as identified in (B). **(E)** 3’UTR shortened genes (red) and lengthened genes (blue) in NIH3T3-HRAS^G12V^ cells as compared to NIH3T3-empty cells. **(F)** USCS genome browser track showing reads for *Timp2* in NIH3T3-empty and NIH3T3-HRAS^G12V^ cells. **(G)** Northern blotting result based on a probe targeting *Timp2* ORF region. Both long and short *Timp2* isoforms are detected in the same blot. A total of 10 µg of RNA was loaded into each lane. **(H)** Western blotting result showing RAS and the secreted Timp2 protein levels. α-Tubulin was used as a loading control.

We next extracted total cellular RNA from HRAS^G12V^-expressing and control cells, and carried out 3’ end sequencing by using the QuantSeq method (see Materials and Methods for detail, **Fig. 1A)**. Each cDNA fragment was sequenced by the paired-end method, yielding a forward read and a reverse read. The former was used for differential gene expression (DGE) analysis, while the latter for APA isoform analysis (see Materials and Methods for detail).

With two biological replicates, we identified 829 up-regulated genes and 879 down-regulated genes **(Figs. 1B and S1E)**. The data from the two replicates were well-corelated (*r* = 0.58, Pearson correlation; **Fig. S1F**). Gene ontology (GO) analysis indicated that top biological processes associated with upregulated genes included “anatomical structure morphogenesis”, “cell adhesion and cell migration” **(Fig. 1C)**, whereas “external encapsulating structure organization” and “regulation of cell population proliferation” were top terms associated with down-regulated genes **(Fig. 1D)**. These GO terms are in agreement with the phenotypes we observed with HRAS^G12V^ cells **(Figs. S1B-S1D)**, i.e., a slower growth rate and a higher migratory ability compared to non-transformed cells.

We next carried out APA isoform analysis^34^ (see Materials and Methods for detail). We identified 40 genes with significant 3’UTR lengthening and 75 genes with 3’UTR shortening in HRAS^G12V^-transformed cells compared to the control (**Fig. 1E**. However, GO analysis did not reveal any biological processes significantly enriched for either gene, indicating that 3’UTR regulation is largely uncoupled from gene expression changes.

Requiring substantial changes in both gene expression and 3’UTR size, we found that the *Timp2* gene, encoding tissue inhibitor of metalloproteinase 2, displayed both 3’UTR shortening (among the top 5 based on the p-value) and expression downregulation in HRAS^G12V^-transformed cells **(Fig. 1F)**. *Timp2* expresses two major APA isoforms **(Fig. S1G)**, with the short isoform containing a 3’UTR of ∼110 nt and the long isoform a 3’UTR of ∼2.6 kb. As such, these two isoforms differ by ∼2.5 kb in the 3’UTR, a region named alternative 3’UTR (aUTR). This is quite remarkable, given that the 5’UTR and coding sequence (CDS) of *Timp2* gene are 433 and 660 bases, respectively. Therefore, the 3’UTR size of long isoform accounts for 70% of the entire transcript, whereas only 9% of the short isoform corresponds to 3’UTR. Notably, both proximal and distal PASs are highly conserved across mammals^35^, This unique transcript configuration difference between isoforms suggests that the aUTR of *Timp2* may have some important functions.

By Northern blot analysis, we verified the expression of the two APA isoforms of *Timp2* in NIH3T3 cells **(Fig. 1G)**. Interestingly, the long isoform had a lower expression level in HRAS^G12V^-transformed cells compared to non-transformed control cells **(Fig. 1G)**. By contrast, the expression level of *Timp2* short isoform had no discernable difference between the cells **(Fig. 1G)**. This result suggests that downregulation of *Timp2* gene expression in cell transformation is attributable mainly to decreased abundance of long isoform.

To examine if *Timp2* isoforms differ in RNA stability in HRAS^G12V^-transformed vs. non-transformed cells, we carried out transcriptional shutdown by Actinomycin D (ActD, **Fig. S1H**, followed by Northern blot analysis of the two isoforms over time. We found no difference in RNA stability between the two isoforms; both isoforms are quite stable, with stability levels similar to that of *Gapdh* transcripts^36,37^ (**Fig. S1H**).

Because Timp2 is a secreted protein, we next examined its protein abundance in cell culture media. By Western blotting, we found that the secreted Timp2 from HRAS^G12V^-transformed cells had a significantly lower abundance compared to that of control **(Fig. 1H)**, indicating decreased expression of secreted Timp2 protein. Since only the long isoform of *Timp2* was decreased in expression in HRAS^G12V^-transformed cells, we conclude that downregulation of *Timp2* long isoform is responsible for decreased Timp2 protein secreted from transformed NIH3T3 cells.

### *Timp2* long APA isoform is the major contributor of Timp2 secreted protein

To investigate the role of *Timp2* aUTR in NIH3T3 cells, we generated two cell lines that express small hairpin RNAs (shRNAs) targeting the open reading frame region of *Timp2* (*sh*ORF) or the aUTR region (*sh*aUTR, **Fig. S2A**), respectively. As expected, only *Timp2* long isoform was knocked down in the *sh*aUTR cell line, while both the long and short isoforms were knocked down in the *sh*ORF cell line **(Figs. 2A & 2B)**. We then examined the secreted Timp2 protein level in culture medium and total cell lysate. As expected, compared to the control cell line (*sh*Ctrl), both *Timp2* aUTR and ORF KD cells had significantly decreased levels of secreted Timp2 protein **(Figs. 2C & 2D)**. Interestingly, *Timp2* aUTR and ORF KD cells showed similar levels secreted protein, suggesting that long isoform KD is chiefly responsible for the secreted protein level change in ORF KD cells. In other words, the *Timp2* long isoform is the major contributor of secreted Timp2 protein **(Fig. 2C & 2D)**.

**Figure 2.**
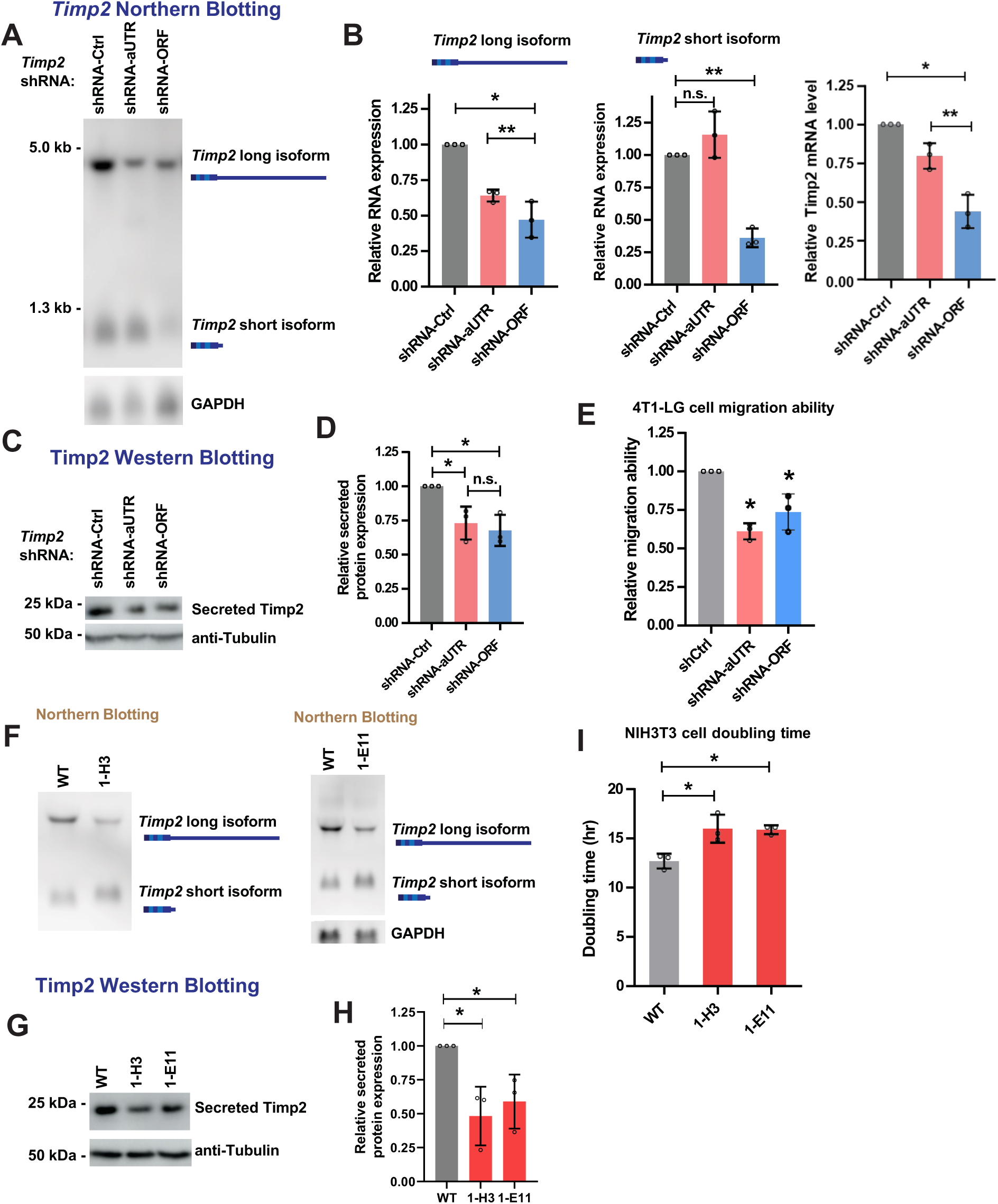
Characterization of *Timp2* shRNA KD and CRISPR/CAS9 aUTR KO^+/-^ cell lines. **(A)** Northern blotting result showing *Timp2* isoform levels in NIH3T3 expressing shRNA-Ctrl, shRNA-aUTR (KD of only the *Timp2* long mRNA isoform) or shRNA-ORF (KD of both *Timp2* short and long mRNA isoforms). **(B)** Quantification of (A). **(C)** Western blotting result showing secreted Timp2 protein level of NIH3T3 cells expressing shRNA-Ctrl, shRNA-aUTR or shRNA-ORF. **(D)** Quantification of (C). **(E)** Migration assay results of 4T1 cells expressing shRNA-Ctrl, shRNA-aUTR or shRNA-ORF. **(F)** Northern blotting result showing *Timp2* isoforms in two NIH3T3 *Timp2* aUTR^+/-^ cell lines, named 1-H3 and 1-E11. **(G)** Western blotting result showing secreted Timp2 protein level (F). **(H)** Quantification of (G). **(I)** Cell doubling time of 1-H3 and 1-E11 compared to NIH3T3 wild type (WT) cells.

We next wanted to examine how *Timp2* KD by *sh*ORF or *sh*aUTR impacts cell migration. Because of the low migration rate of NIH3T3 cells at the baseline **(Fig. S2D)**, we carried out *Timp2* aUTR and ORF KDs in mouse 4T1 cells, a commonly used mouse breast cancer metastasis model which has a high migration rate. Consistent with the NIH3T3 cell data, we observed decreased *Timp2* mRNA expression and the amount of secreted protein in 4T1 cells with aUTR and ORF KDs **(Figs. S2B &S2C)**. Using the wound healing migration assay (see Materials and Methods for detail), we found that both *Timp2* aUTR and ORF KD cells displayed a slower cell migration rate compared to control cells **(Fig. 2E)**.

We next set out to remove the aUTR sequence of *Timp2* in NIH3T3 cell lines by using the CRISPR/Cas9 system. We designed a pair of Cas9-guide RNAs targeting a position right after the *Timp2* proximal PAS (pPAS) and a position ∼200 nt upstream of the distal PAS (dPAS) **(Fig. S2D)**. A pair of PCR primers targeting the start and end of *Timp2* long 3’UTR were used for genotyping after CRISPR/Cas9 **(Fig. S2D)**. Two heterozygous knockout (KO) cell lines of *Timp2* aUTR (1-H3 and 1-E11) were obtained, which were validated by both genotyping PCR and Sanger sequencing **(Fig. S2E)**. Notably, despite multiple tries, we were not able to obtain homozygous KO cells, suggesting that *Timp2* aUTR might be essential for cell survival or growth. Northern blotting analysis indicated that both 1-H3 and 1-E11 aUTR-KO^+/-^ cell lines had significantly decreased expression of *Timp2* long isoform and a lower *Timp2* long isoform to short isoform expression ratio **(Fig. 2F)**. In addition, similar to *Timp2 sh*aUTR KD cells, the secreted Timp2 protein level was down-regulated in the two *Timp2* aUTR KO^+/-^ cell lines **(Fig. 2G & 2H)**. Moreover, both 1-H3 and 1-E11 cell lines had slower growth rates compared to control cells transfected with a random gRNA **(Figure 2I)**. Taken together, analyses of *Timp2* aUTR KD and KO cells support the notion that the *Timp2* long isoform is the major contributor of secreted Timp2 protein.

### *Timp2* aUTR KO^+/-^ cells and HRAS^G12V^-transformed cells show a commonly regulated gene set

To further explore the functional importance of *Timp2* aUTR, we extracted total RNA from the two *Timp2* aUTR KO^+/-^ cell clones (1-H3 and 1-E11) and carried out RNA sequencing using the QuantSeq method. We identified 353 genes that were commonly up-regulated in the two cell clones and 400 genes that were commonly down-regulated **(Figs. 3A-3C, and Figs. S3A & S3B)**. GO analysis indicated that gene associated with “cytoskeleton organization”, “movement of cell or subcellular component”, and “cell adhesion” tended to be up-regulated in the two *Timp2* aUTR KO^+/-^ cell clones **(Fig. 3D)**. It is noteworthy that many of the GO terms were also enriched for regulated genes in HRAS^G12V^-transformed cells **(Fig. 1C & 1D)**.

**Figure 3.**
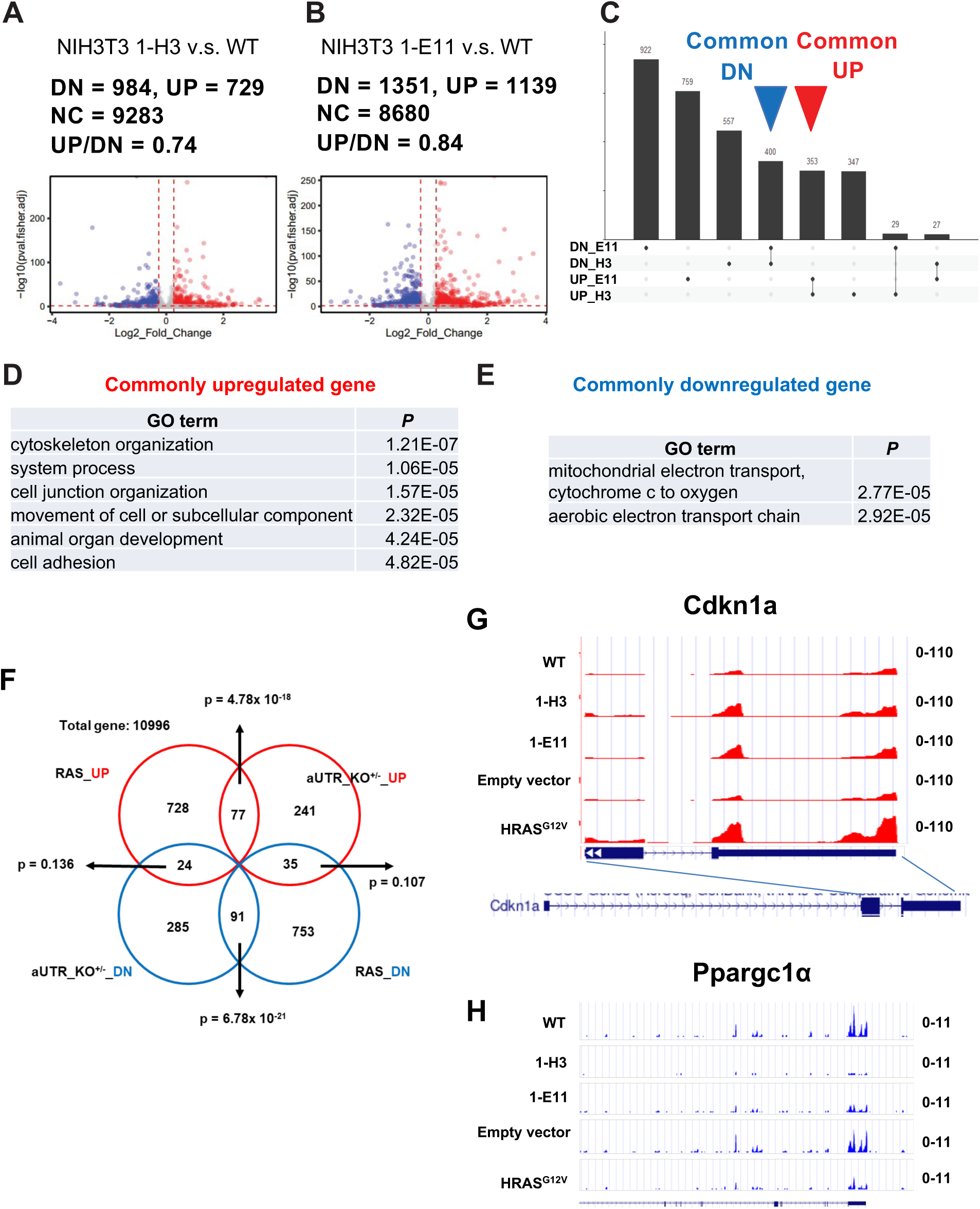
HRAS^G12V^ transformation and *Timp2* aUTR knockout (KO) lead to similar gene expression changes. **(A)** Upregulated and downregulated genes (*P* < 0.05, Fisher’s exact test) in 1-H3 cells compared to WT. **(B)** Upregulated and downregulated genes (*P* < 0.05, Fisher’s exact test) in 1-E11 cells compared to WT. **(C)** UpSet plot showing commonly upregulated and downregulated genes in (A) and (B). **(D)** Top GO terms enriched for commonly upregulated genes in 1-H3 and 1-E11 cells. **(E)** Top GO terms enriched for commonly downregulated genes in 1-H3 and 1-E11 cells. **(F)** Fisher’s exact test of paired group of genes: (1) upregulated genes in HRAS^G12V^ cells, (2) downregulated genes in HRAS^G12V^ cells, (3) upregulated genes in *Timp2* aUTR KO^+/-^ cells, and (4) downregulated genes in *Timp2* aUTR KO^+/-^ cells. P-value was calculated for each intersection. **(G)** USCS genome browser tracks showing reads mapped to *Cdkn1a* in five cell lines. **(H)** USCS genome browser tracks showing reads mapped to *Ppargc1α* in five cell lines.

Interestingly, several GO terms related to mitochondrial functions were enriched for commonly down-regulated genes **(Fig. 3E)**. To verify this, we measured the respiration rate of *Timp2* aUTR KO^+/-^ cells, and found that both the basal respiration rate and the ATP-linked respiration rate were significantly down-regulated in the *Timp2* aUTR KO^+/-^ cells as compared to control cells **(Fig. S3C & S3D)**. Notably, decreased mitochondrial respiration rate was also observed in HRAS^G12V^-transformed NIH3T3 cells compared to control cells **(Figs. S3E & S3F)**.

We next directly compared gene expression changes in aUTR KO^+/-^ and HRAS^G12V^-transformed cells (**Figs. 1 and 3)**. We found a significant overlap between these two cell types, with 77 up-regulated genes shared by HRAS^G12V^ cells and *Timp2* aUTR KO^+/-^ cells (p = 4.78 x 10^-18^) and 91 down-regulated genes (p = 6.78 x 10^-21^) shared by both. This result suggests that down-regulation of the *Timp2* long isoform might be responsible for a sizable portion of genes regulated during cell transformation by HRAS^G12V^ **(Fig. 3F)**. The full lists of these shared genes are shown in **Figs. S3G & S3H**; two example genes, *Cdkn1a* and *Ppargc1α*, are shown in **Figs 3G and 3H**. Note that these two genes are key regulators of cell proliferation and mitochondrial functions, respectively (see Discussion), whose downregulation could play key roles in the cell phenotypes we have observed.

### Down-regulation of *Timp2* long isoform mitigates gene expression regulation by HRAS^G12V^

NIH3T3 cells with constitutive overexpression of HRAS^G12V^ represents a stable, long-term cell transformation model. To gain temporal information about gene regulation by HRAS^G12V^, we generated an NIH3T3 cell line transduced with a lentiviral vector containing HRAS^G12V^ under the control of a tetracycline response element (TRE)-containing promoter (see Materials and Methods for detail). As such, HRAS^G12V^ expression can be induced by doxycycline (Dox)**(Figs. 4A & S4A).** For simplicity, these cells were named NIH3T3/iHRAS^G12V^ cells. We carried out RNA sequencing with cells treated with Dox (Dox+) for 48 hrs and carried our transcriptomic profiling by QuantSeq, using cells without Dox treatment (Dox-) as a control. We found an overall positive correlation in gene expression changes (*r* = 0.48, Pearson correlation; **Fig. S4E**) between NIH3T3/iHRAS^G12V^ cells and cells with constitutive HRAS^G12V^ expression **(Figs. 4B & 4C)**. We identified 667 common up-regulated genes and 709 common down-regulated while only 11 genes showed opposite expression changes. This result indicates that most of the gene expression changes caused by HRAS^G12V^ transformation might be established within 48 hrs of its expression.

**Figure 4.**
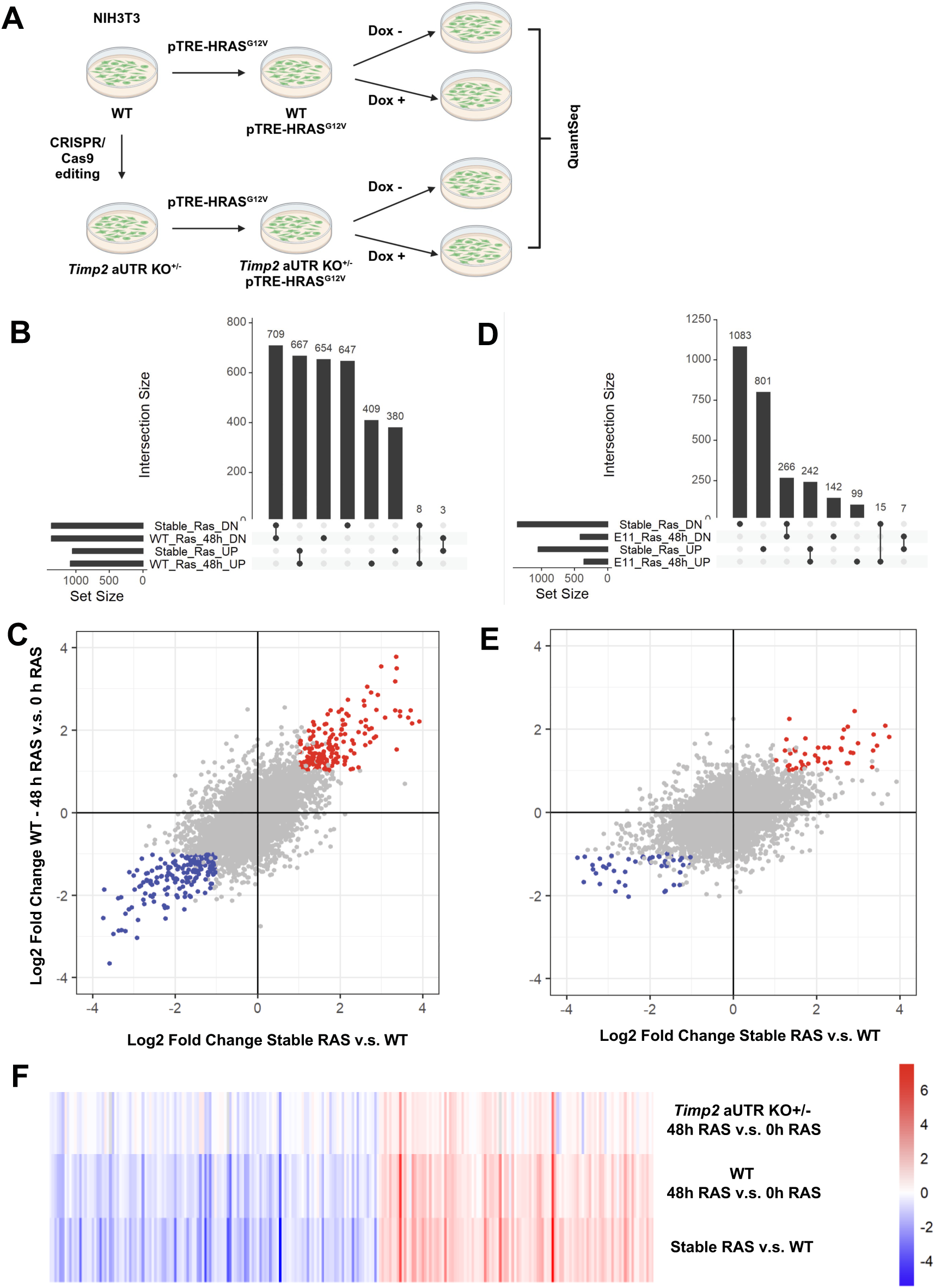
Preemptive down-regulation of *Timp2* long isoform mitigates gene expression changes by further HRAS^G12V^ activation. **(A)** Diagram showing generation of inducible HRAS^G12V^ cell lines based on NIH3T3 WT or NIH3T3 aUTR KO^+/-^ cells. Induction of HRAS^G12V^ expression was carried out Dox treatment for 48 hours. RNA samples were subjected to RNA seqeuencing based on the QuantSeq-Pool method. **(B)** UpSet plot showing commonly regulated genes between (1) stable HRAS^G12V^ vs. WT cells and (2) Dox+ vs. Dox-treatment of aUTR KO^+/-^ cells containing iHRAS^G12V^. **(C)** Scatter plot of commonly regulated genes identified in (B). **(D)** UpSet plot showing commonly regulated genes between (1) stable HRAS^G12V^ vs WT cells and (2) Dox+ vs. Dox-treatment of aUTR KO^+/-^ cells containing iHRAS^G12V^. **(E)** Scatter plot of commonly regulated genes identified in (D). **(F)** Heatmap of genes with expression changes in three comparisons. The intensity of color indicates the extent of regulation.

Because of the substantial overlap of regulated genes between cells with HRAS^G12V^ transformation and *Timp2* aUTR KO^+/-^ cells, we hypothesized that Timp2 regulation might be an effector event of HRAS^G12V^ signaling. To test this hypothesis, we generated a *Timp2* aUTR KO^+/-^ cell line with Dox-inducible HRAS^G12V^ **(Fig. 4A & S4B)**, and then compared the differentially expressed genes after induction of HRAS^G12V^ for 48 hrs in aUTR KO^+/-^ cells vs. wild-type cells. Interestingly, while the trend of gene regulation by HRAS^G12V^ in aUTR KO+/- cells is similar to that in wild-type cells, the expression fold changes overall were smaller in aUTR KO+/- cells as compared to wild-type cells **(Figs 4D & 4E)**. This result indicates that down-regulation of *Timp2* long isoform, as in *Timp2* aUTR KO^+/-^ cells, could mitigate gene expression changes elicited by HRAS^G12V^ **(Fig. 4F)**, supporting the notion that downregulation of *Timp2* long isoform is a key signaling event activated by HRAS^G12V^.

We can also compare the regulated genes using the two pairs of the Dox+ / Dox-samples **(Figure S4C-4D, S4G)**. Overall, the trend of gene regulation in aUTR KO+/- background looks similar when compared to either the stable RAS cell line **(Figure 4E)** or the inducible RAS cell line **(Figure S4D)**. Actin-related genes were enriched in the GO terms analysis of Timp2 aUTR KO^+/-^ cells versus WT cells, HRAS^G12V^ transformed cells versus empty cells, and inducible HRAS^G12V^ cells. To verify this, we performed beta-actin immunofluorescence assay on the cells mentioned above (**Figure S4H-S4J**). Compared to the control cells, all the cells with HRAS^G12V^ expression or Timp2 aUTR KO^+/-^ undergoes changes in morphology.

### Distinct mRNA localization features of short and long 3’UTR isoforms of *Timp2*

The differential contributions of *Timp2* long vs. short isoforms to secreted protein levels motivated us to further explore functions of *Timp2* aUTR. We reasoned that *Timp2* aUTR might play a role in mRNA localization to the endoplasmic reticulum (ER), where translation of mRNAs encoding secreted proteins takes place.

To examine *Timp2* mRNA localization in NIH3T3 cells, we carried out single molecule fluorescence in situ hybridization (smFISH)^38^. We designed two sets of smFISH probes, one targeting the common region to both isoforms (*Timp2*_P) and the other the aUTR sequence, i.e., specific to the long isoform (*Timp2*_D, **Fig. 5A**). These two probe sets were labelled with two fluorescent dyes, namely, Texas Red and Quasar 670. After hybridization and imaging, we first analyzed smFISH dots identified by the two fluorescent channels, each of which represents one mRNA molecule. We reasoned that an overlap between smFISH dots in the two channels would support the existence of the long isoform (set 1), whereas a short isoform would give rise to a dot in the channel only for the common region probe set (set 2) **(Fig. 5A)**. In addition, after cell segmentation, we identified the centroid positions of set 1 and set 2 dots in each cell and calculated the average distance of each set of dots to its centroid position, aiming to capture overall distributions of the two isoforms from the center of each cell. To quantitatively represent the relative distribution of isoforms, we used the ratio of average distances of two dot sets (set 1 average distance / set 2 average distance, **Fig. S5A**), which was named relative distribution index (RDI). As such, the two isoforms have similar subcellular distributions around the centroid if the RDI = 1. As a proof-of-concept for RDI measurement, we designed smFISH probes for *Tfrc* mRNA, which is known to be localized to the ER, and *Gapdh* mRNA, which is known to be diffusive in the cytosol ^39^. We labelled *Tfrc* probes with Texas Red and *GAPDH* probes with Quasar 670, and repeated the experiment with dye swap **(Fig. S5B & S5C)**. In both experiments, we observed a significant difference in subcellular localization of the two probe sets and, consistently, RDI = 0.74 in all the NIH3T3 cell lines tested **(Fig. S5D)**. This result indicates that RDI for smFISH signals could distinguish mRNA localization.

**Figure 5.**
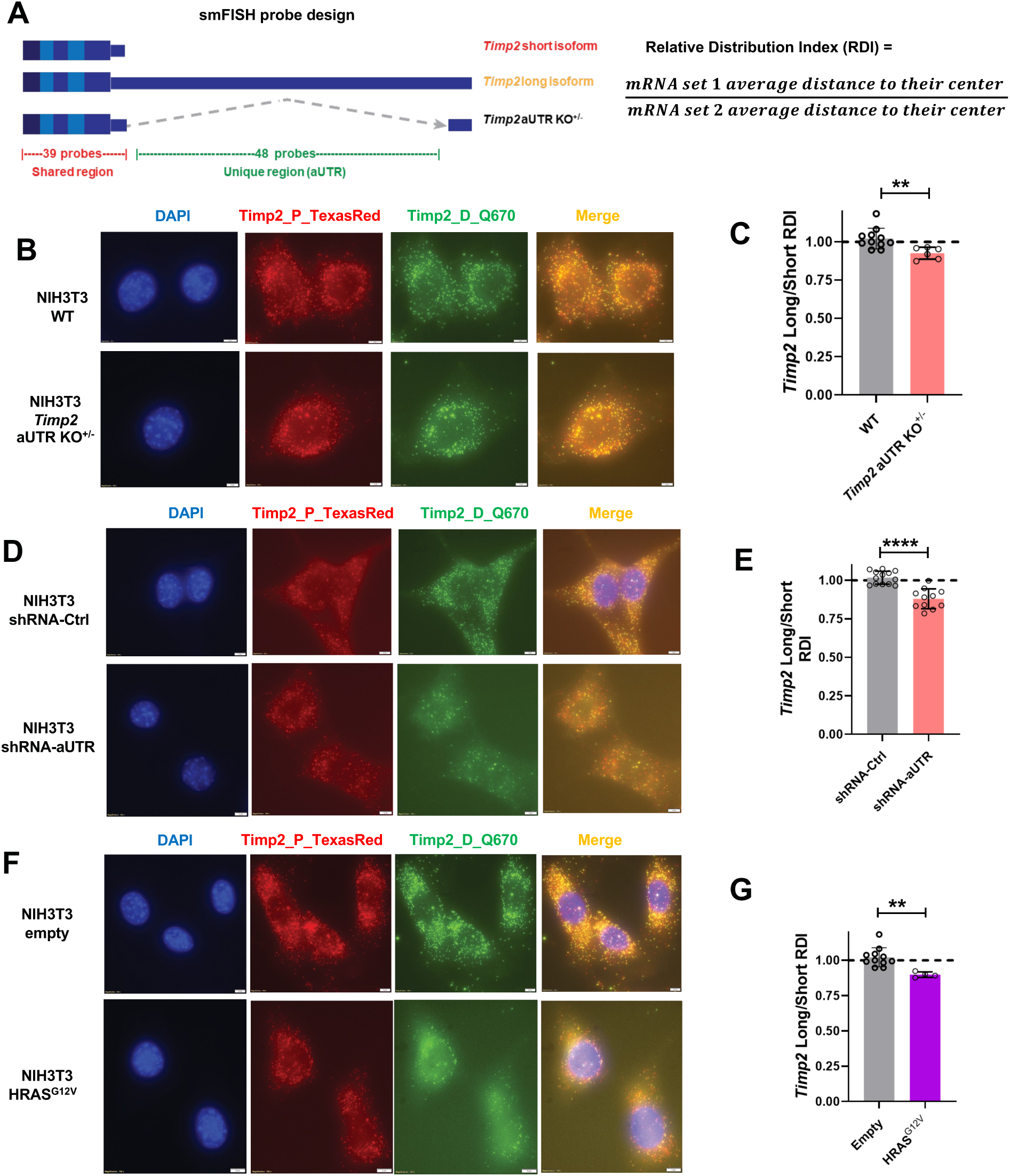
smFISH analysis of *Timp2* short and long isoforms. **(A)** Diagram showing the design of two sets of smFISH probes for *Timp2*. **(B)** Example smFISH data of NIH3T3-Timp2-aUTR KD and control cells. Pseudo colors are shown. **(C)** Quantification of (B). **(D)** Example smFISH data of NIH3T3-Timp2-aUTR KO^+/-^ and control cells. Pseudo colors are shown. **(E)** Quantification of (D). **(F)** Example smFISH data of NIH3T3-HRAS^G12V^ and control cells. Pseudo colors are shown. **(G)** Quantification of (F).

We next carried out smFISH for *Timp2* long and short isoforms in *Timp2 sh*aUTR knockdown cells, *Timp2* aUTR KO^+/-^ cells and HRAS^G12V^ cells as well as their respective control cells (indicated as wild type cells, **Figs. 5B-5G**). The three control cell lines had RDI = 1, indicating no discernable localization differences betwen the two *Timp2* short and long isoforms. Interestingly, in all the three cell lines where there was a lower ratio of *Timp2* long / short isoform compared to wild type cells, RDI< 1 was obtained, indicating that *Timp2* long isoform is closer to the cell centroid or perinuclear space, where the ER is enriched, as compared to the short isoforms in these cells **(Fig. 5B-5G)**. These data suggest a connection between *Timp2* aUTR-mediated mRNA subcellular localization and the level of long 3’UTR isoform.

To corroborate the smFISH result, we fractionated cells into cytosol and membrane portions by using the sequential detergent wash method ^40^(**Fig. S5E;** see Materials and Methods for detail). RNA from the cytosol and the membrane fractions were extracted and subjected to RNA-seq by the QuantSeq method. To measure the relative distribution of each transcript between membrane and cytosol fractions, we calculated a membrane localization score (MLS) based on the ratio of abundance of each pA site-defined transcript (using reads per million mapped [RPM]) between the fractions ^41^ **(Fig. S5E;** see Materials and Methods for detail). We found that both *Timp2* isoforms were enriched in the membrane fraction, presumably the ER (**Fig. S5F**), consistent with the fact that Timp2 is a secreted protein. However, the long isoform had a higher MLS value than the short isoform (2.32 vs. 2.04, **Fig. S5F**), indicating that the former has a greater membrane association than the latter. Interestingly, in cells transformed by HRAS^G12V^, both long and short isoforms showed decreased MLS (1.90 and 1.42, respectively, **Fig. S5F).** But the difference between the two isoforms, or ΔMLS (long isoform – short isoform) was greater in transformed cells than wild type cells (ΔMLS = 0.48 vs. 0.28, **Fig. S5F)**. Strikingly, in aUTR KO^+/-^ cells, while the short isoforms (combining the wild type short isoform and the isoform with aUTR removed) had a lower MLS compared to wild type cells (MLS = 1.26), the long isoform had increased MLS (2.92). As such, the difference between the two isoforms was the greatest in these cells (ΔMLS = 1.66). Together, these data are in good agreement with smFISH data, supporting the notion that the *Timp2* long 3’UTR isoform has a higher potential to be membrane-associated than the short 3’UTR isoform, and this difference is enhanced in transformed cells.

### Biased subcellular localization of *Timp2* long 3’UTR isoform correlates with translational efficiency and protein secretion

To address whether the biased localization of *Timp2* long 3’UTR isoforms has any impact on the abundance of secreted protein, we performed smFISH in wild-type cells and *Timp2* aUTR KO^+/-^ cells with or without inhibition of mRNA translation by puromycin treatment **(Figure 6A)**. In wild-type cells, RDI = 1 regardless of translational inhibition. By contrast, the differential localization of the long and short isoforms in *Timp2* aUTR KO^+/-^ cells was observed in cells without translational inhibition (RDI <1) but disappeared after puromycin treatment (RDI = 1) **(Figure 6B)**, indicating that the biased subcellular localization of *Timp2* long 3’UTR isoform in *Timp2* aUTR KO^+/-^ cells is dependent on protein translation.

**Figure 6.**
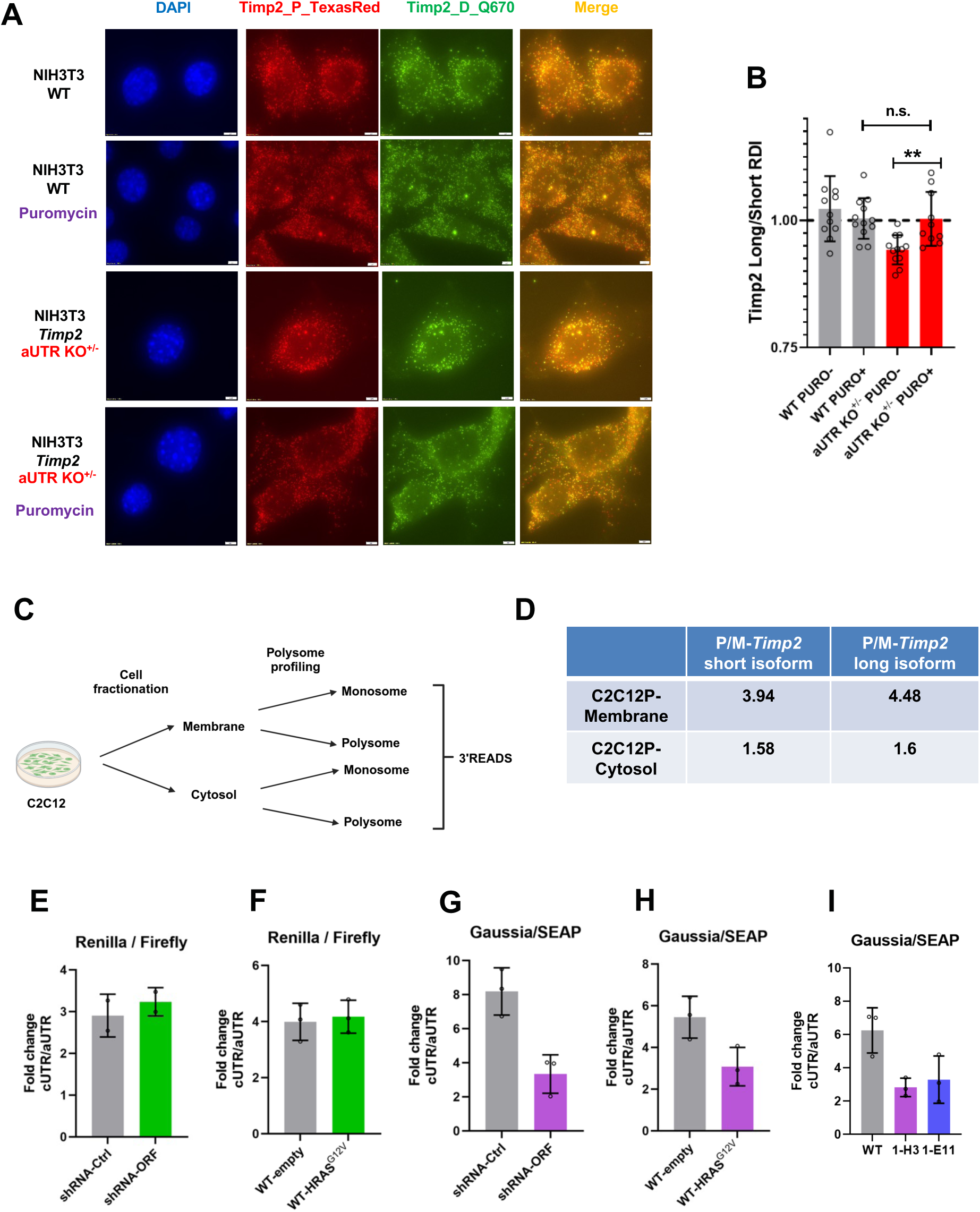
Biased subcellular distribution of *Timp2* long isoform in aUTR KO ^+/-^ cells correlates with mRNA translation on the ER and protein secretion. **(A)** smFISH data of *Timp2* isoforms in NIH3T3 WT and *Timp2* aUTR KO^+/-^ cells with or without puromycin treatment. Pseudo colors are shown. **(B)** Quantification of the four groups of samples in (A). **(C)** Diagram showing cell fractionation followed by polysome profiling and 3’READS+ sequencing of each fraction in C2C12 cells. **(D)** UCSC genome tracks showing *Timp2* short and long isoform peaks measured in (C). **(E)** Renilla and firefly luciferase assays in NIH3T3 cells expressing shRNA-Ctrl or shRNA-ORF. **(F)** Renilla and firefly luciferase assays in NIH3T3-empty and NIH3T3-HRAS^G12V^ cells. **(G)** Gaussia luciferase and SEAP assays in NIH3T3 cells expressing shRNA-Ctrl or shRNA-ORF. **(H)** Gaussia luciferase and SEAP assays in NIH3T3-empty and NIH3T3-HRAS^G12V^ cells. **(I)** Gaussia luciferase and SEAP assays in NIH3T3 WT, 1-H3 and 1-E11 cells.

Polysome profiling can be used to examine translational activity of transcripts ^41^. To this end, we re-analyzed polysome profiling data our lab previously generated with C2C12 cells. The data corresponded to 3’ end sequencing of transcript in monosome and polysome fractions of C2C12 cells in both cytosol and membrane fractions ^41,42^ **(Fig. 6C)**. As expected, both the short and long *Timp2* isoforms were substantially enriched in the membrane fraction. However, the P/M value (reflecting transcript abundance in polysome fraction vs. monosome fraction) of *Timp2* long isoform was significantly higher than that of the *Timp2* short isoform in the membrane fraction **(Fig. 6D)**, indicating that the aUTR of *Timp2* has an enhancing role for ER-dependent mRNA translation of the *Timp2* long isoform.

To corroborate the impact of *Timp2* aUTR on gene expression, we next constructed two series of vectors for reporter assays. In the first series, the cUTR and aUTR sequences of *Timp2* were cloned into a vector (pRF) containing Renilla and firefly luciferase sequences. UTR sequences were placed after the *Renilla* luciferase coding region, and the firefly luciferase was used as an internal control **(Fig. S6A)**. Both Renilla and Firefly luciferases are non-secreted proteins that remain within the cell after translation. In the second series, UTR sequences were cloned into a vector (pGS) containing *Guassia* luciferase and secreted alkaline phosphatase (SEAP) sequences. The UTRs were placed after the *Gaussia* luciferase and SEAP was used as an internal control **(Fig. S6B)**. Both *Gaussia* luciferase and SEAP are secreted proteins.

We first expressed pRF vectors in HRAS^G12V^-transformed cells and *Timp2 sh*ORF KD cells as well as their matching control cells **(Fig. S6C)**. We compared the pRF-cUTR / pRF-aUTR expression levels in different cell lines and found that this ratio remained the same across all cell lines tested **(Fig. 6E-6F)**. In contrast, the pGS-cUTR / pGS-aUTR expression levels decreased in HRAS^G12V^, *Timp2 sh*ORF, and aUTR KO^+/-^ cells **(Figs. 6G-6I)**. Taken together, these results indicate that *Timp2* aUTR had a positive effect on the production of secreted proteins but not intracellular proteins.

In summary, as illustrated in **Fig. 7**, our results indicate that *Timp2* 3’UTR shortening is an effector event of RAS signaling. The aUTR of *Timp2* functions to enrich long *Timp2* isoform on the ER and enhances mRNA translation. The selective downregulation of long 3’UTR isoform in HRAS^G12V^-transformed cells leads to decreased levels of secreted Timp2 protein, which further elicits gene expression changes, leading to inhibited cell proliferation and enhanced cell migration.

**Figure 7.**
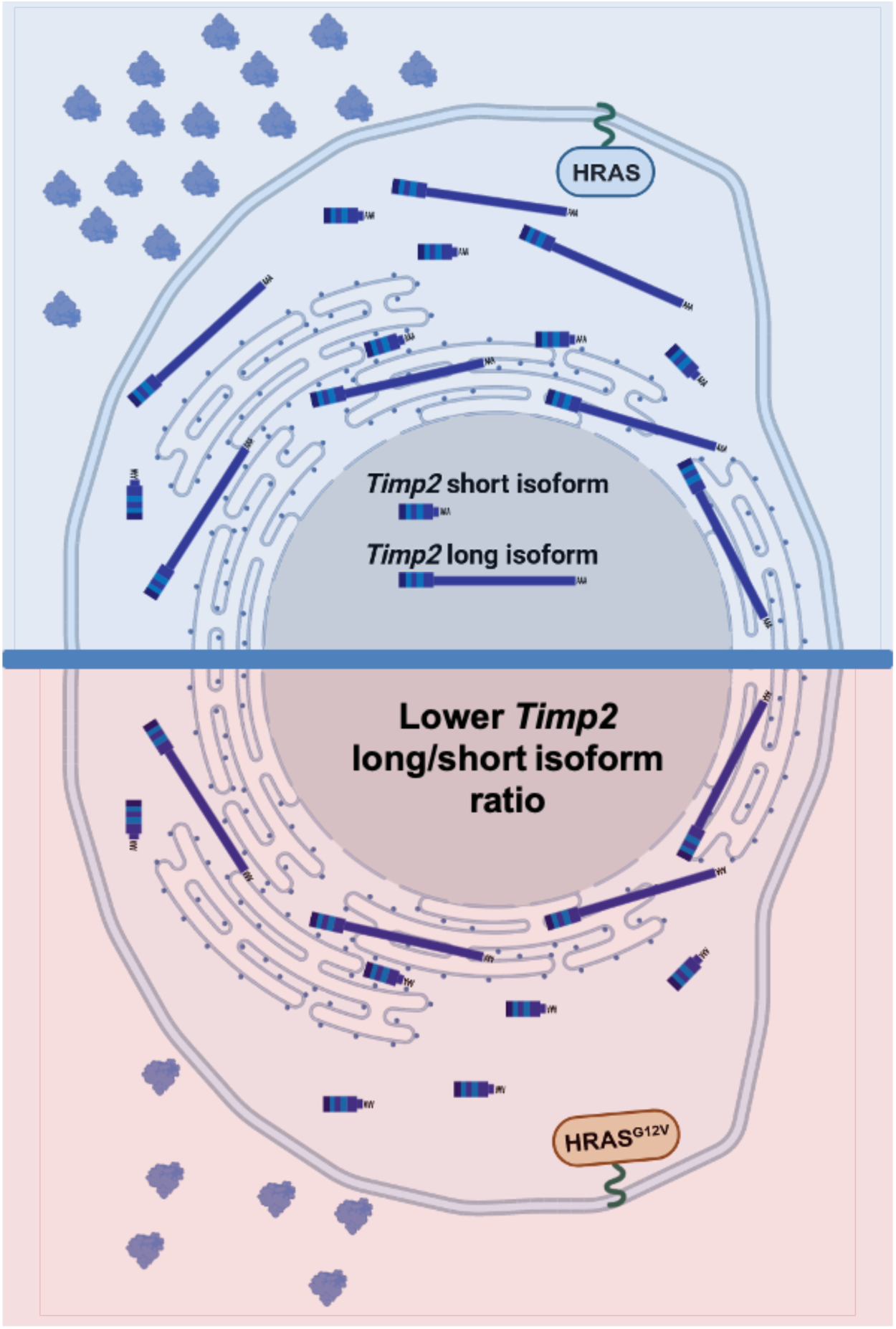
Schematic summarizing regulation of *Timp2* short and long 3’UTR isoforms in wild type or RAS-transformed NIH3T3 cells.

## DISCUSSION

In this study, we identified *Timp2* 3’UTR shortening as an effector event of HRAS^G12V^-mediated NIH3T3 transformation. We explored the subcellular localization differences between the two 3’UTR isoforms as well as their contributions to Timp2 protein expression.The human *TIMP2* and mouse *Timp2* genes have been well studied for more than three decades since their discovery ^43^, including their downregulation in RAS-transformed cells ^44,45^. However, none of the prior work has examined the importance of their 3’UTR isoforms. The current study indicates that regulation of *Timp2* 3’UTR isoform expression is an integral part of cell transformation by HRAS^G12V^.

While we show that HRAS activation leads to decreased expression of *Timp2* long 3’UTR isoform with the short 3’UTR isoform largely unchanged, the underlying mechanism is unclear. Our lab previously used 4-thio-uridine (4sU)-based metabolic labeling of RNA to examine isoform stability differences in several human and mouse cell lines, including human HEK293T, HepG2, and SH-SY5Y cells ^46^ and mouse NIH3T3 and C2C12 cells ^41^. Re-analysis of these data indicates that the short and long 3’UTR isoforms do not differ significantly in mRNA stability (data not shown). In addition, a previous study by Hammani et al. using transcriptional shutdown with Actinomycin D and Northern blotting showed that *TIMP2* short and long isoforms had similar RNA stability levels in human fibrosarcoma cell line HT1080 ^47^. Interestingly, the authors also found that *TIMP2* transcripts had comparable stability levels to β-actin mRNA, which is generally considered to be quite stable ^48^. We also performed similar Actinomycin D-based mRNA decay assays in NIH3T3 cells and found both the long and the short Timp2 mRNAs are very stable (Fig. S1H). Therefore, the two 3’UTR isoforms of *Timp2* do not appear to differ in half-lives in different cell types and conditions.

We posit that downregulation of long 3’UTR isoform might be due to inhibition of the usage of distal pA site in transformed cells. Because we did not observe global APA changes genome-wide, i.e, similar numbers of genes showing 3’UTR shortening vs. those showing 3’UTR lengthening, the mechanism for Timp2 APA regulation could involve specific regulation of the distal pA site. This needs to be further explored in the future.

We show that the aUTR sequence of *Timp2* impacts subcellular localization. One plausible mechanism is that some sequence motifs in the aUTR may interact with cognate RNA-binding proteins (RBPs) for localization to the ER. Notably, there are several islands of conserved subregions in *Timp2* aUTR, especially in the second half ^9^. Future studies using deletion mutations or blocking RBP binding, such as using dCas13-based systems^49^, could help identify functional elements responsible for the subcellular localization functions of *Timp2* aUTR. In addition, it is possible that the sheer size of *Timp2* aUTR may have an impact on its mRNA localization. On this note, we recently revealed a translation-independent ER association mechanism, or TiERA, for many transcripts ^41^. Transcript size, GC content, and RNA structures were found to be important for TiERA. It would be interesting to examine how the TiERA model can explain the difference in ER association between the two Timp2 isoforms and, more importantly, whether RAS transformation could alter the TiERA mechanism leading to different ER association potentials for the same transcript ^41^.

TIMP2 was originally found to bind matrix metalloproteinase 2 (MMP2) on the cell surface ^50^ and to inhibit its metalloproteinase activity ^43^. Later reports indicated that TIMP2 can also activate MMP2 as well as other MMPs ^51^. Not surprisingly, TIMP2/Timp2 have been reported to both promote and inhibit cell proliferation ^52–55^ but others indicating inhibition of cell proliferation ^56–59^. In fact, the effect of Timp2 on cell growth can be quite different in even closely related melanoma cell lines ^60^. Our data showing that Timp2 promotes cell proliferation of NIH3T3 cells and cell migration of 4T1 cells are in line with the notion that Timp2’s function is cell context-specific. Our lab has previously identified *Timp2* as one of the top genes with significant 3’UTR lengthening in differentiation of mouse C2C12 myoblast cells ^9^. We previously found that many of the APA events regulated in C2C12 differentiation are also regulated in other cell differentiation systems, such as neurogenesis ^46^. It is plausible that *Timp2* APA correlates with global APA changes in cell differentiation and development. One open question is whether *Timp2* APA is also regulated by RAS-dependent signaling in these cell differentiation systems. Notwithstanding the underlying mechanisms, the fact that *Timp2* APA isoforms are dynamically regulated in cell differentiation and transformation points to an important connection between APA regulation and Timp2-mediated regulation of cell proliferation, migration, and extracellular matrix remodeling.

## STAR★METHODS

### KEY RESOURCES TABLE

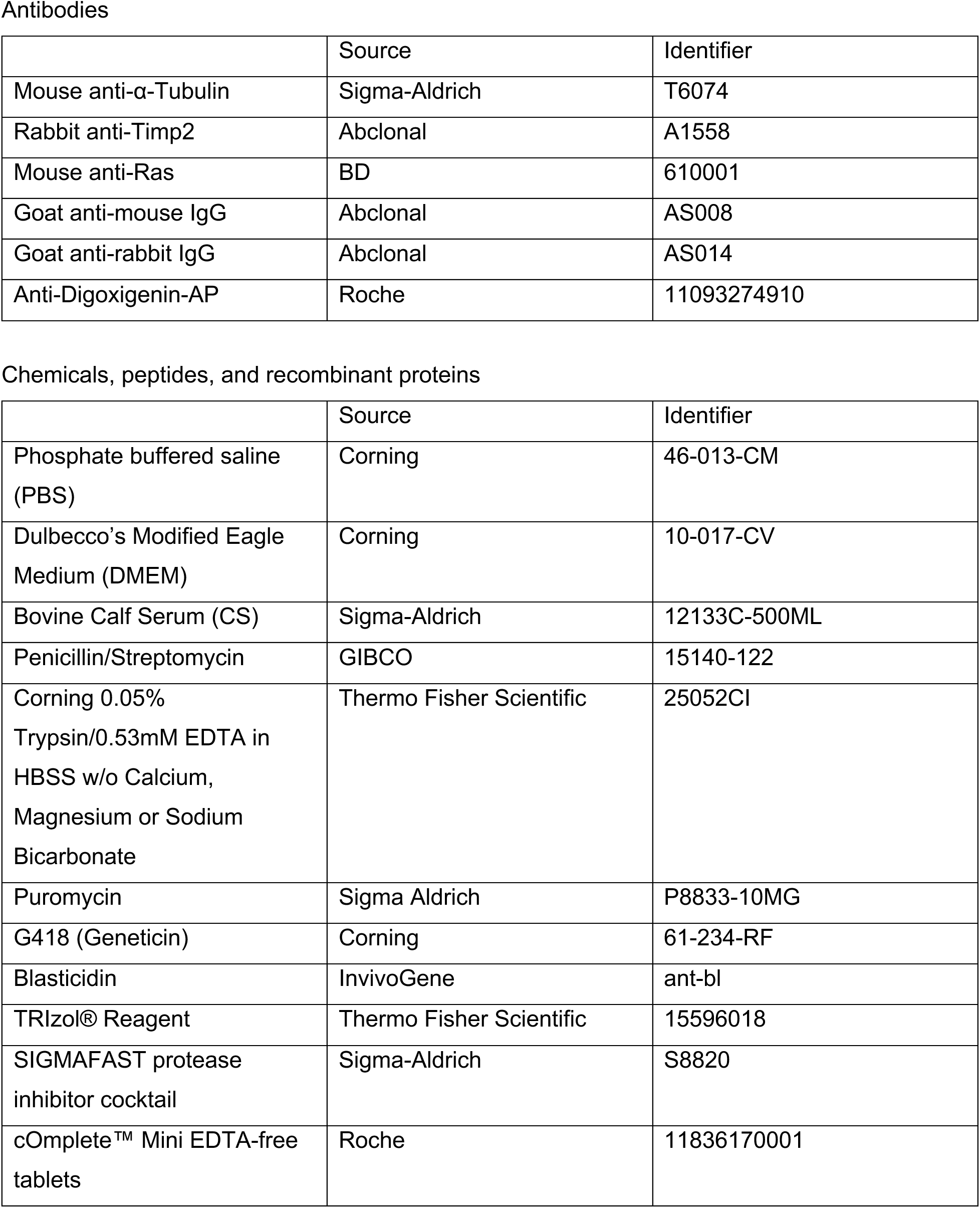
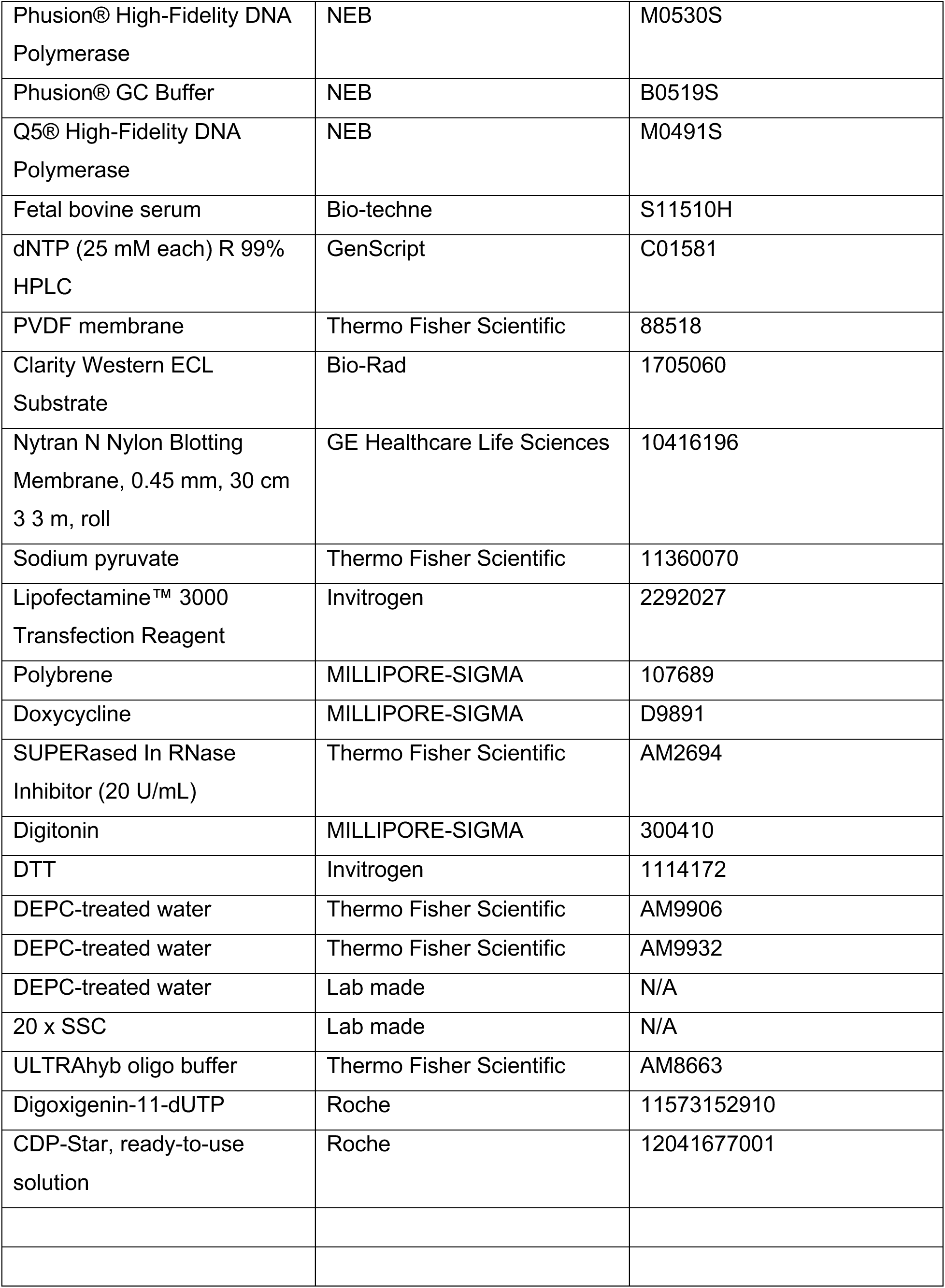
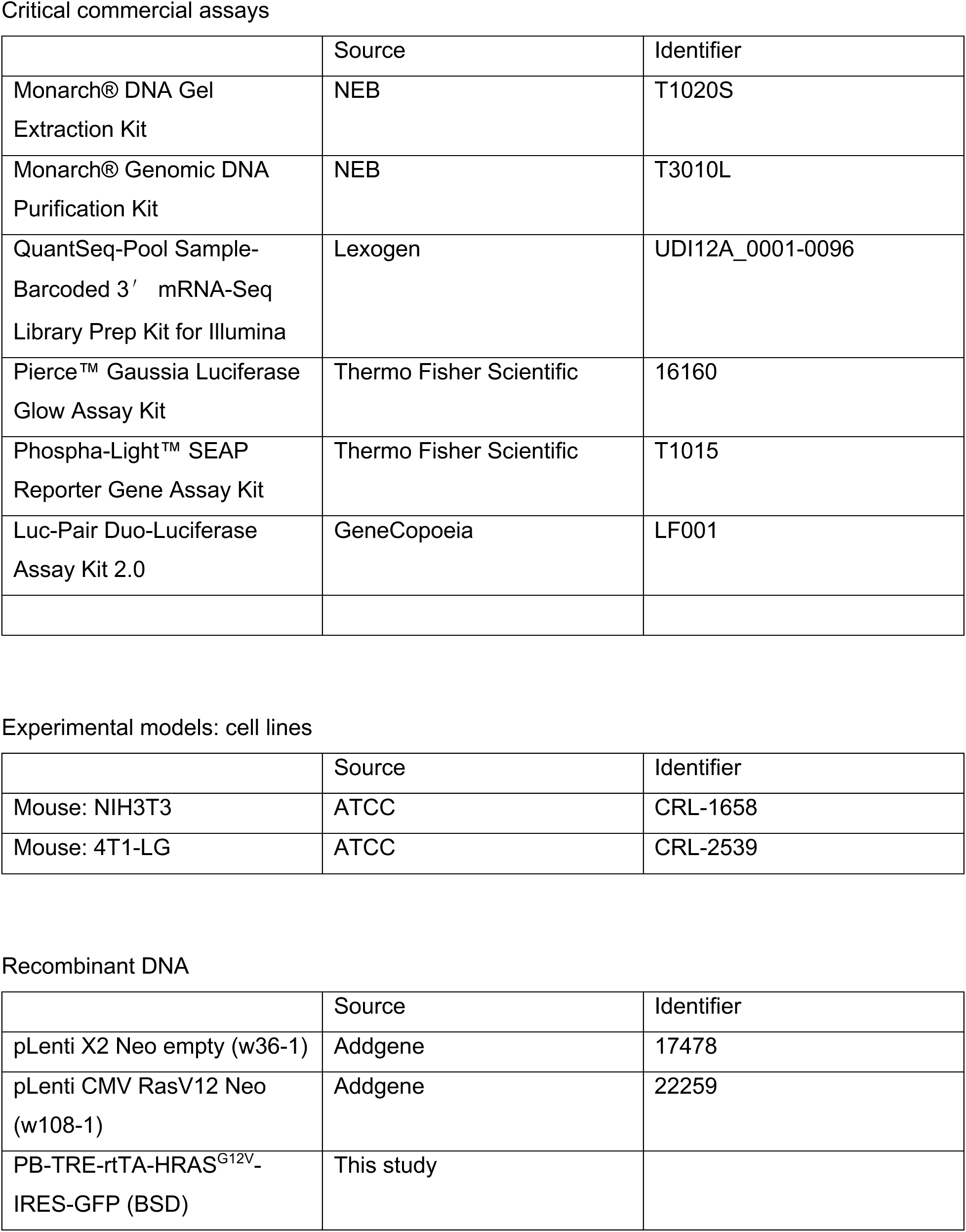
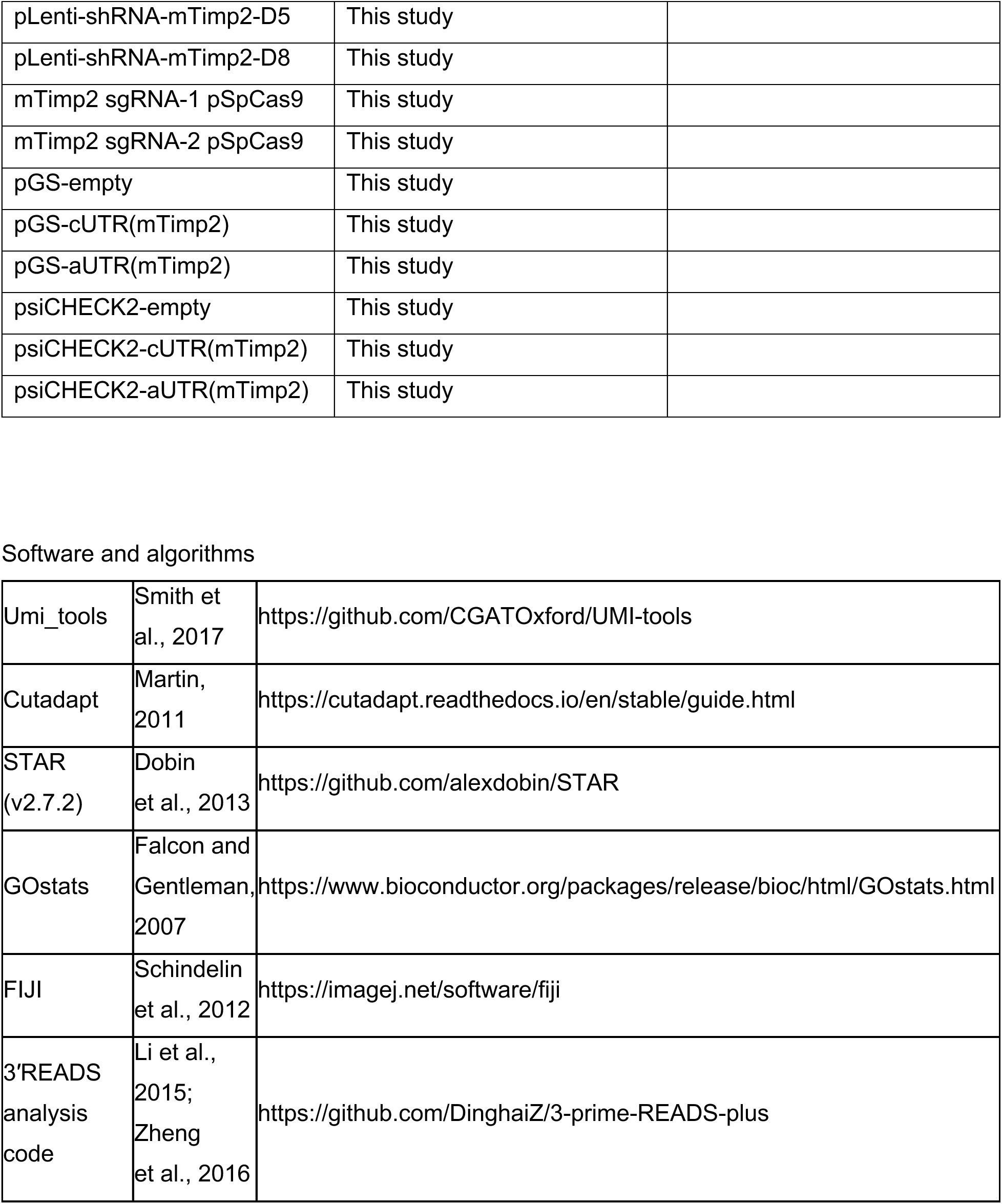

Antibodies
|  | Source | Identifier |
| --- | --- | --- |
| Mouse anti-α-Tubulin | Sigma-Aldrich | T6074 |
| Rabbit anti-Timp2 | Abclonal | A1558 |
| Mouse anti-Ras | BD | 610001 |
| Goat anti-mouse IgG | Abclonal | AS008 |
| Goat anti-rabbit IgG | Abclonal | AS014 |
| Anti-Digoxigenin-AP | Roche | 11093274910 |

Chemicals, peptides, and recombinant proteins
|  | Source | Identifier |
| --- | --- | --- |
| Phosphate buffered saline (PBS) | Corning | 46-013-CM |
| Dulbecco's Modified Eagle Medium (DMEM) | Corning | 10-017-CV |
| Bovine Calf Serum (CS) | Sigma-Aldrich | 12133C-500ML |
| Penicillin/Streptomycin | GIBCO | 15140-122 |
| Corning 0.05% Trypsin/0.53mM EDTA in HBSS w/o Calcium, Magnesium or Sodium Bicarbonate | Thermo Fisher Scientific | 25052CI |
| Puromycin | Sigma Aldrich | P8833-10MG |
| G418 (Geneticin) | Corning | 61-234-RF |
| Blasticidin | InvivoGene | ant-bl |
| TRIzol® Reagent | Thermo Fisher Scientific | 15596018 |
| SIGMAFAST protease inhibitor cocktail | Sigma-Aldrich | S8820 |
| cOmplete™ Mini EDTA-free tablets | Roche | 11836170001 |
| Phusion® High-Fidelity DNA Polymerase | NEB | M0530S |
| Phusion® GC Buffer | NEB | B0519S |
| Q5® High-Fidelity DNA Polymerase | NEB | M0491S |
| Fetal bovine serum | Bio-techne | S11510H |
| dNTP (25 mM each) R 99% HPLC | GenScript | C01581 |
| PVDF membrane | Thermo Fisher Scientific | 88518 |
| Clarity Western ECL Substrate | Bio-Rad | 1705060 |
| Nytran N Nylon Blotting Membrane, 0.45 mm, 30 cm 3 3 m, roll | GE Healthcare Life Sciences | 10416196 |
| Sodium pyruvate | Thermo Fisher Scientific | 11360070 |
| Lipofectamine™ 3000 Transfection Reagent | Invitrogen | 2292027 |
| Polybrene | MILLIPORE-SIGMA | 107689 |
| Doxycycline | MILLIPORE-SIGMA | D9891 |
| SUPERased In RNase Inhibitor (20 U/mL) | Thermo Fisher Scientific | AM2694 |
| Digitonin | MILLIPORE-SIGMA | 300410 |
| DTT | Invitrogen | 1114172 |
| DEPC-treated water | Thermo Fisher Scientific | AM9906 |
| DEPC-treated water | Thermo Fisher Scientific | AM9932 |
| DEPC-treated water | Lab made | N/A |
| 20 x SSC | Lab made | N/A |
| ULTRAhyb oligo buffer | Thermo Fisher Scientific | AM8663 |
| Digoxigenin-11-dUTP | Roche | 11573152910 |
| CDP-Star, ready-to-use solution | Roche | 12041677001 |

Critical commercial assays
|  | Source | Identifier |
| --- | --- | --- |
| Monarch® DNA Gel Extraction Kit | NEB | T1020S |
| Monarch® Genomic DNA Purification Kit | NEB | T3010L |
| QuantSeq-Pool Sample-Barcoded 3' mRNA-Seq Library Prep Kit for Illumina | Lexogen | UDI12A_0001-0096 |
| Pierce™ Gaussia Luciferase Glow Assay Kit | Thermo Fisher Scientific | 16160 |
| Phospha-Light™ SEAP Reporter Gene Assay Kit | Thermo Fisher Scientific | T1015 |
| Luc-Pair Duo-Luciferase Assay Kit 2.0 | GeneCopoeia | LF001 |

Experimental models: cell lines
|  | Source | Identifier |
| --- | --- | --- |
| Mouse: NIH3T3 | ATCC | CRL-1658 |
| Mouse: 4T1-LG | ATCC | CRL-2539 |

Recombinant DNA
|  | Source | Identifier |
| --- | --- | --- |
| pLenti X2 Neo empty (w36-1) | Addgene | 17478 |
| pLenti CMV RasV12 Neo (w108-1) | Addgene | 22259 |
| PB-TRE-rtTA-HRAS<sup>G12V</sup>-IRES-GFP (BSD) | This study |  |
| pLenti-shRNA-mTimp2-D5 | This study |  |
| pLenti-shRNA-mTimp2-D8 | This study |  |
| mTimp2 sgRNA-1 pSpCas9 | This study |  |
| mTimp2 sgRNA-2 pSpCas9 | This study |  |
| pGS-empty | This study |  |
| pGS-cUTR(mTimp2) | This study |  |
| pGS-aUTR(mTimp2) | This study |  |
| psiCHECK2-empty | This study |  |
| psiCHECK2-cUTR(mTimp2) | This study |  |
| psiCHECK2-aUTR(mTimp2) | This study |  |
| Software and algorithms |  |  |
| Umi_tools | Smith et al., 2017 | https://github.com/CGATOxford/UMI-tools |
| Cutadapt | Martin, 2011 | https://cutadapt.readthedocs.io/en/stable/guide.html |
| STAR (v2.7.2) | Dobin et al., 2013 | https://github.com/alexdobin/STAR |
| GOstats | Falcon and Gentleman, 2007 | https://www.bioconductor.org/packages/release/bioc/html/GOstats.html |
| FIJI | Schindelin et al., 2012 | https://imagej.net/software/fiji |
| 3'READS analysis code | Li et al., 2015; Zheng et al., 2016 | https://github.com/DinghaiZ/3-prime-READS-plus |

## EXPERIMENTAL MODEL AND STUDY PARTICIPANT DETAILS

### Cell culture

HEK293T cells were cultured at 37°C and 5% CO_2_ in Dulbecco’s modified Eagle’s medium (DMEM) containing high glucose (Corning, 10-017-CV) supplemented with 10% (v/v) fetal bovine serum (FBS, Bio-techne, S11510H) and 1% (v/v) penicillin-streptomycin (GIBCO, 15140-122).

NIH3T3 cells were cultured at 37°C and 5% CO_2_ in DMEM containing high glucose supplemented with 10% (v/v) calf serum (Sigma, 12133C), 1% (v/v) penicillin-streptomycin and 1% (v/v) 100mM sodium pyruvate (Thermo Fisher Scientific, 11360070).

4T1-luc2-GFP cells (containing GFP and luciferase genes, PerkinElmer) were cultured at 37°C and 5% CO_2_ in RPMI 1640 medium with 10% (v/v) FBS and 1% (v/v) penicillin-streptomycin.

### Generation of vectors

#### HRAS^G12V^ vectors

pLenti CMV RasV12 Neo (Addgene # 22259) was used to generate NIH3T3-HRAS^G12V^ cells and pLenti X2 Neo empty (Addgene # 17478) was used to generate NIH3T3-empty cells.

#### Timp2 shRNA vectors

Mouse *Timp2* shRNA vectors were ordered from Wistar Institute TRC Cloning Vector Stock with the following product numbers:

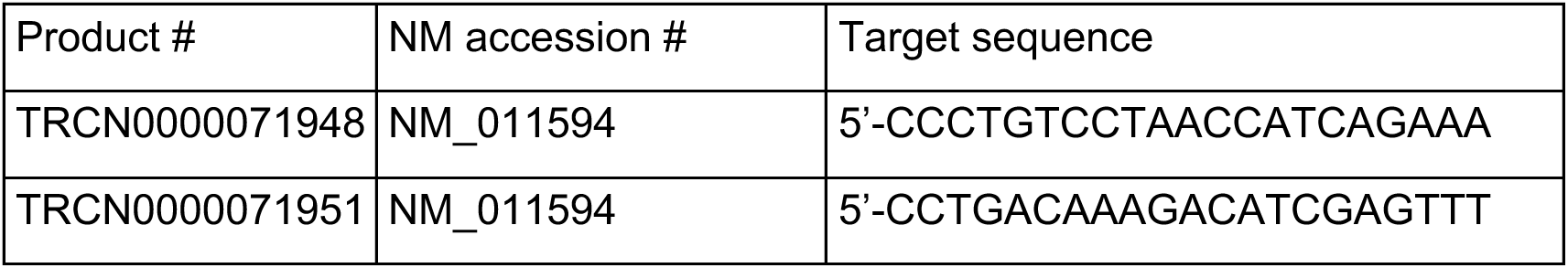

#### Timp2 aUTR CRISPR/Cas9 vectors

DNA fragments containing mTimp2 sgRNA-1 or -2 with sticky overhangs were inserted into pSpCas9(BB)-2A-Puro (PX459) (Addgene# 48139) by using the Bbs I site. The sequences of sgRNAs are as follows:

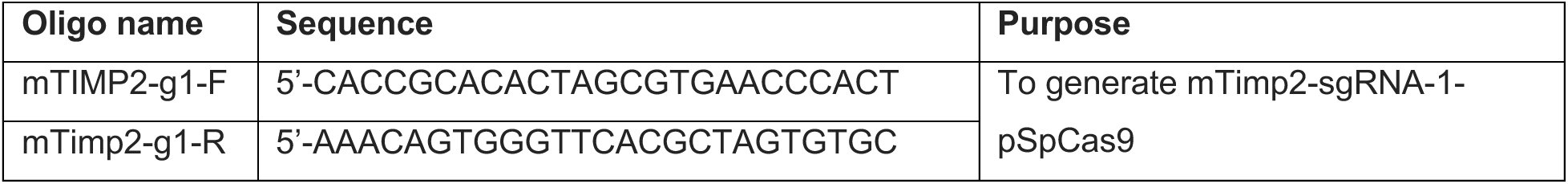
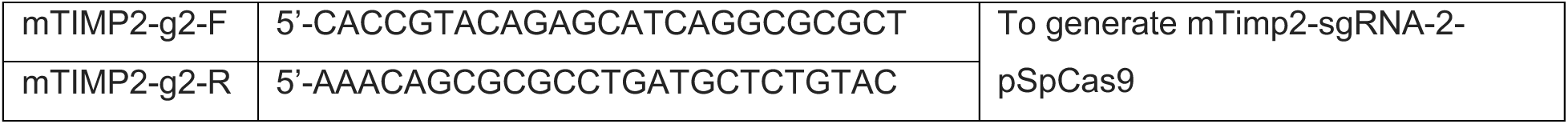

#### Inducible HRAS^G12V^ vectors

##### PB-TRE-rtTA-HRAS^G12V^-IRES-GFP

HRAS^G12V^ was PCR-amplified with the following primers and the template pLenti CMV RasV12 Neo (w108-1) (Addgene# 22259). PCR products were cut by Nhe I and BamH I, and were inserted into pCW-3FLAG-FIP1-V1-IRES-EGFP-BSD ^34^ with the same enzymes to generate pCW-HRAS^G12V^-IRES-EGFP-BSD.

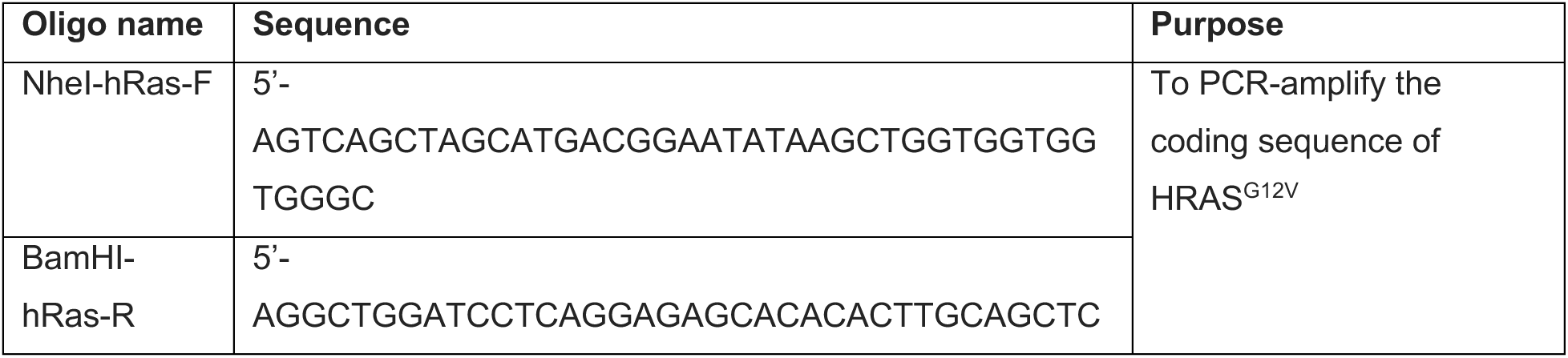

#### Reporter assay vectors

For Renilla / firefly reporter assay vectors, mouse Timp2 3’UTR was amplified from genomic DNA (obtained from NIH3T3) and inserted into psiCHECK2 (Catalog # C8021, Promega) by using Xho I and Not I to generate psiCHECK2-cUTR(mTimp2), aUTR(mTimp2), and FullUTR(mTimp2).

For Gaussia / SEAP reporter assay vectors, Gaussia and SEAP coding sequences were PCR-amplified from pCMV-Gaussia-Dura Luc (Catalog # 16191, ThermoFisher) and pSEAP2-Control (Catalog # 631717, Takara). The NEBuilder method (Catalog # E2621S, NEB) was used to swap Gaussia and SEAP sequences with Renilla and firefly luciferase sequences in psiCHECK2, respectively, to generate pGS versions. With the same method, mTimp2 UTR was inserted into the pGS vector.

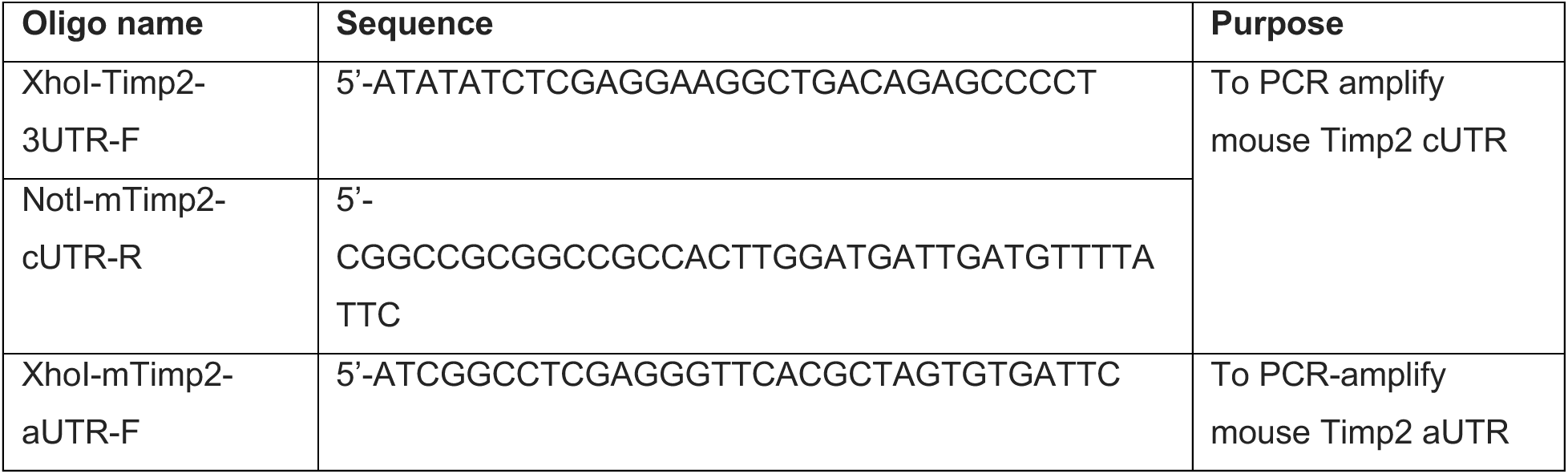
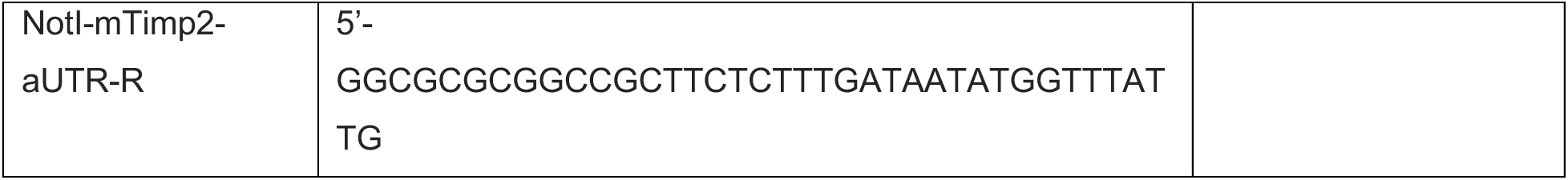

### Generation of *Timp2* aUTR knockout cells using CRISPR/Cas9

NIH3T3 cells were seeded on a 12-well plate at 1 x 10^5^ cells per well. A mixture of 500 ng mTimp2 sgRNA-1 pSpCas9 and 500 ng mTimp2 sgRNA-2 pSpCas9 vectors was transfected into 1 x 10^5^ cells by using Lipofectamine 3000 (Catalog # 2292027, Invitrogen) according to manufacturer’s instructions. Cell culture media was changed 6 hrs post transfection, followed by cell culture for another 48 hrs. Cells were then subject to puromycin (1.5 µg/mL) selection for 7 days. Selected cells were resuspended in DMEM + 0.5% FBS at a concentration of 4 x 10^6^ / mL and single cell sorting was performed by using the MoFlo AstriosEQ cell sorter (Beckman Coulter). Sorted cells in 96-well plates were kept under normal cell culture condition for 5 days and well-grown colonies were selected under the microscope, labelled, and amplified to reach an appropriate number of cells. Genomic DNA was then extracted from each cell clone by using the Monarch® Genomic DNA Purification Kit (Catalog # T3010L, NEB) and was used for genotyping PCR with the following protocol for each DNA sample: each PCR reaction contained 0.5 µL Genomic DNA (50-150 ng / μL), 0.5 µL 10 μM forward primer (5’-ATATATCTCGAGGAAGGCTGACAGAGCCCCT), 0.5 µL 10 μM reverse primer (5’-ATATATGCGGCCGCCAACTGAGGCACACCTTCAG), 6.2 µL double distilled water, 2 µL 5x Phusion® GC Buffer (NEB, B0519S), 0.1 µL Phusion® High-Fidelity DNA Polymerase (NEB, M0530S), and 0.2 µL 10 mM dNTPs (Catalog # C01581, GenScript). The PCR reaction involves initiation (98 °C, 30 sec), 30 cycles of amplification (98°C 8 sec and then 72 °C 1 min 40 sec), extension (72 °C, 7 min). The samples generated two bands at ∼2,750 bp and ∼320 bp were identified as Timp2 aUTR KO^+/-^ cell lines. The low molecular weight bands were cut from the gel and purified with the Monarch® DNA Gel Extraction Kit (Catalog # T1020S, NEB), and sent for Sanger sequencing to verify engineered genomic sequences around the *Timp2* aUTR.

### Generation of stable cell lines using lentivirus

HEK293T cells were grown to 90% confluency on a 6-well plate. Each well was used to generate one lentivirus construct. Each well was transfected with a mixture containing 2 µg vector of interest, 1 µg psPAX2 vector and 1 µg pMD2.G vector. Transfection was carried out by using the Lipofectamine 3000 (Catalog # 2292027, Invitrogen) according to the manufacturer’s instructions. Media was changed after 4-6 hrs. Cells were then incubated for another 48 hrs before supernatant (6 mL per well) was harvested at 24 hrs, 36 hrs and 48 hrs post transfection. Media was changed each time supernatant was harvested.

#### Timp2 shRNA knock-down cell lines

Media containing shRNA-Control, shRNA-Timp2-aUTR and shRNA-Timp2-ORF lentivirus was collected as mentioned above. NIH3T3 cells or 4T1-luc2-GFP cells were seeded on a 12-well plate at 1 x 10^5^ cells per well in 0.5 mL media. Lentivirus-containing medium (0.5 mL) was added to each well, together with 0.8 µL of 10 mg/mL polybrene (Catalog # 107689, Sigma) to reach a total volume of 1 mL per well. After cell culture for another ∼12 hrs, media was changed. Cells were then grown for another 24-36 hrs before 2 ug/mL puromycin (Sigma, P8833) was added for selection. After 5-7 days of selection, cells were collected as Timp2 shRNA knock-down cell lines.

#### HRAS^G12V^ inducible cells

Cell cuture media containing pCW-rtTA-HRAS^G12V^-IRES-GFP lentivirus was generated and collected as mentioned above. NIH3T3 cells with or without CRISPR/Cas9 editing were seeded on a 12-well plate at 1 x 10^5^ cells per well in 0.5 mL media. Lentivirus-containing medium (0.5 mL) was added to each well together with 0.8 µL of 10 mg/mL polybrene to reach a total volume of 1 mL per well. After cell culture for another ∼12 hrs, media was changed. Cells were then grown for another 24-36 hrs before 20 ug/mL Blasticidin (Catalog code: ant-bl, InvivoGene) was added for selection. After 5-7 days of selection, HRAS^G12V^ inducible cell lines were collected.

### Mammalian expression plasmids, transfections, and RNA isolation

NIH3T3 cells were seeded on 6-well plates and grown to 80%-90% confluency at the time of RNA isolation. For doxycycline (Dox)-inducible cell lines, 2 µg/mL Dox (Catalog # D9891, Sigma) was added to induce the expression of HRAS^G12V^ or hTIMP2. Total RNA was isolated 40-48 hrs later by using TRIzol (Catalog # 15596018, Thermo Fisher Scientific) according to manufacturer’s instructions.

### Cell Fractionation

We followed the cell fractionation protocol based on sequential detergent extraction as previously developed by Jagannathan et al. ^40^ with minor modifications. Briefly, NIH3T3 cells on 10-cm dishes were rinsed with pre-chilled PBS and incubated in pre-chilled PBS with 1 mg/mL MgCl_2_ on ice for 20 min. After removing the PBS/MgCl_2_ buffer completely, 0.75 mL permeabilization buffer (110 mM KCl, 25 mM K-HEPES pH 7.4, 2.5 mM MgCl_2_, 0.1 mM EGTA, 0.015% (w/v) digitonin (Catalog # 300410, Sigma), 1 mM DTT, 1 x FAST Protease Inhibitor Cocktail (Catalog # S8820, Sigma), and 40 U/mL SuperaseIn RNase Inhibitor (Catalog # AM2694, Thermo Fisher Scientific) was added to each dish to cover all the cells evenly. After incubation on ice for 10 min, the dishes were tilted on ice to allow collection of the cytosolic fraction. The dishes were then gently washed with 8 mL pre-chilled wash buffer (110 mM KCl, 25 mM K-HEPES pH 7.4, 2.5 mM MgCl_2_, 0.1 mM EGTA, 0.004% (w/v) digitonin, 1 mM DTT [Catalog # 1114172, Invitrogen], 1 x FAST Protease Inhibitor Cocktail, and 40 U/mL SuperaseIn RNase Inhibitor). After removing all the wash buffer, 0.75 mL lysis buffer (200 mM KCl, 25 mM K-HEPES pH 7.4, 10 mM MgCl_2_, 1% (v/v) NP40, 0.5% (w/v) sodium deoxycholate, 1 mM DTT, 1 x FAST Protease Inhibitor Cocktail, and 40 U/mL SuperaseIn RNase Inhibitor) was added to each dish to cover all the cells evenly. After incubation on ice for 10 min, the dishes were tilted on ice for collection of the membrane fraction. The dishes were then gently washed with 8 mL pre-chilled PBS buffer. After removing all the PBS, the insoluble fraction on the dish was saved at -80 °C before RNA extraction. The cytosolic and membrane fractions were centrifuged at 700 x g and subsequently 1,400 x g for 5 min in a micro-centrifuge to remove any debris.

### QuantSeq FWD and QuantSeq-Pool mRNA library preparation and sequencing

RNA samples were diluted in DEPC-treated water (Catalog # AM9906, Thermo Fisher Scientific). Qubit and TapeStation were used to measure the concentration and quality of the RNA samples, respectively. RNA samples with an RNA Integrity Number (RIN) = 10 were used for cDNA library preparation. QuantSeq FWD or QuantSeq-Pool mRNA library preparation was performed according to manufacturer’s instruction (Lexogen). After library preparation, each library pool was subject to QC and was sequenced on an Illumina Hiseq or NovaSeq system at Admera Health.

### Differential gene expression analysis

QuantSeq-Pool data were processed according to manufacturer’s recommendation. Briefly, For QuantSeq FWD data, read 1 data were first trimmed by using umi_tools ^61^ and cutadapt ^62^, followed by mapping to the mouse genome (mm9) by using STAR-2.7.7a ^63^. The number of reads mapped to each gene was counted by using featureCounts ^64^. For QuantSeq-Pool data, raw data were first de-multiplexed by using the idemux tool, and read 1 data were then used. The Umi_tools package was used to remove read duplicates based on the location of the alignment and the unique molecular identifier (UMI) information. The number of reads mapped to each gene was calculated by using the featureCounts tool ^64^. Only genes with more than five reads in a sample were used for further analysis. The read count of each gene was normalized to the total mapped reads to the genome in a sample. PseudoCount of 1 was applied to prevent infinity values in ratio calculation. The significance of expression difference was assessed by the Fisher’s exact test. All *P* values were adjusted by the Benjamini–Hochberg (BH) method to control the false discovery rate (FDR)^65^. BH-adjusted *P* value < 0.05 was considered significant. A fold change of 1.2 or 2 was additionally applied to select regulated genes.

### APA analysis

For QuantSeq-Pool data, read 2 data were used after removal of UMI and poly(dT) sequences. Reads with a mapping quality score (MAPQ) < 10 were discarded. The last aligned position (LAP) of each read was compared to annotated PASs in the polyA_DB database ^35^, allowing ± 24 nt flexibility. Matched reads were poly(A) site-supporting (PASS) reads and were used for further APA analysis. For 3′ UTR APA analysis, the two PASs in the 3′UTR of the last exon with the highest expression levels were compared. One was named proximal PAS (pPAS), and the other distal PAS (dPAS). For IPA analysis, the IPA isoform with the highest expression level among all IPA isoforms was compared to all isoforms using last exon PASs (TPA isoforms) combined. Relative Expression Difference (RED) was calculated as the difference in ratio (log_2_) of isoform abundance (dPAS isoform vs. pPAS isoform) between two comparing samples. Significant APA events were those with RED > log_2_(1.2) or <−log_2_(1.2) and BH-adjusted *P* < 0.05 (Fisher’s exact test).

### Gene Ontology analysis

Gene ontology (GO) analysis was carried out by using the GOstats function in Bioconductor package ^66^. Generic terms (associated with >1,000 genes) were discarded. To reduce redundancy in reporting, a GO term was discarded if its associated genes overlapped with those of another term that had a more significant *P* value by >75%. *P* values are based on the hypergeometric test.

### Northern blotting

Total RNA (2-5 µg per sample) was electrophoresed in a 1.2 % denaturing formaldehyde agarose gel (with a final concentration of 0.6 M formaldehyde) with 1x MOPS running buffer (0.2 M MOPS free acid, 0.05 M Sodium Acetate and Disodium EDTA, pH = 7). The gel was washed twice in nuclease-free water (15 min each) and twice in 10X SSC (15 min each) before RNA transfer to a Nytran^TM^ N Nylon Blotting membrane (Catalog # 10416196, GE Healthcare Life Sciences) overnight. The membrane was then subjected to UV crosslinking in a UV crosslinker (Catalog # FB-UVXL-1000, Fisher Scientific) with 120 mJ of ultraviolet radiation per unit area. The membrane was then incubated in 10 mL of pre-warmed (50 °C) ULTRAhyb oligo buffer (Catalog # AM8663, Thermo Fisher Scientific) in a hybridization tube for 45 min at 42 °C with rotation before 2 µL of probe (described below) was added.

Northern blotting probes for *Timp2* (common to both isoforms only) and *Gapdh* were made by asymmetry PCR with paired PCR primers. Each PCR reaction contained 10 ng vector containing the desired sequence, 1 µL 0.1 μM forward primer (*Gapdh* 5’-TCACCACCATGGAGAAGGC; *Timp2* 5’-TTTCTTGACATCGAGGACCC), 1 µL 10 μM reverse primer (*Gapdh* 5’-GCTAAGCAGTTGGTGGTGCA; *Timp2* 5’-TCCAGGAAGGGATGTCAAAG), 33.18 µL double distilled water, 5 µL 10x Taq Buffer, 0.5 µL Taq DNA Polymerase (M0530, NEB), 0.5 µL 10 mM dATP, 0.5 µL 10 mM dGTP, 0.5 µL 10 mM dCTP, 0.32 µL 10 mM dTTP and 5 µL 1 mM Digoxigenin-11-dUTP (Catalog # 11573152910, Roche). The PCR reaction involved initiation (94 °C, 2 min), 50 cycles of amplification (94°C 20 sec and then 51 °C 30 sec), and extension (72 °C, 2 min).

Membrane was incubated with probes at 42 °C with rotation overnight. Two washes with Low Stringency Wash Solution (2X SSC, 0.1% SDS) were then carried out (5 min each at 42°C), followed by two washes (15 min each at 42°C) with High Stringency Wash Solution (0.1X SSC, 0.1% SDS); washes were carried out with constant agitation. The membrane was then rinsed briefly for 1-5 min in Washing buffer (0.1 M maleic acid, 0.15 M NaCl, pH 7.5 (+20°C), 0.3% (v/v) Tween 20). The membrane was blocked for 30 min in Blocking solution, containing bovine albumin 0.5% (w/v) in Washing buffer, and then incubated at room temperature with antibody solution (anti-Digoxigenin-AP [Roche, 11093274910] diluted 10,000x [75 mU/mL] in 1x Blocking solution) for another 30 min. Membrane was washed twice, 15 minutes each, in Washing buffer. The membrane was equilibrated for 2-5 minutes in Detection buffer (0.1 M Tris-HCl*, 0.1 M NaCl, pH = 9.5). Chemiluminescence reaction was carried out by quickly applying ∼1 mL CDP-Star, ready-to-use solution (Catalog # 12041677001, Roche) to evenly soak the membrane. After a 5 min incubation, chemiluminescent signals were measured by using Amersham Imager 680.

### Western blotting

Cells were lysed 40-48 hrs after seeding or Dox induction by using RIPA buffer (150 mM NaCl, 1% Triton X-100, 50 mM Tris pH 7.5, 0.1% SDS, 0.5% sodium deoxycholate, and protease inhibitors (Catalog # 11836170001, Roche) for 20 min on ice. Cell lysates were then spun at 12,000 x *g* for 15 min and the supernatants were transferred to new tubes. Protein samples were resolved on a 4-12% Bis-Tris gel and transferred to a PVDF membrane (Thermo Fisher Scientific, 88518). Membranes were blocked with 5% nonfat milk for 1 hr before incubation with primary antibody (diluted in 1x TBST) overnight at 4°C. Membranes were then washed with 1x TBST (3 x 5 min) followed by incubation with a secondary antibody at room temperature for 1 hr. The following antibodies were used: mouse anti-α-tubulin (1:2,000, Sigma T6074), rabbit anti-Timp2 (1:1,000, ABclonal, A1558), mouse anti-Ras (1:5,000, BD Transduction Laboratories, 610001), anti-mouse IgG (1:10,000, Abclonal, AS008) and anti-rabbit IgG (1:10,000, Abclonal, AS014). Membranes were subject to chemiluminescent reactions by using the Clarity Western ECL Substrate (Bio-Rad, 1705060) according to the manufacturer’s protocol.

### Immunofluorescence assay

NIH3T3 cells were seeded and cultured on coverslips in 12-well plates at 1 x 10^5^ cells per well for 18-24 hours. To fix samples, cells were washed twice with PBS, fixed with ice-cold 100% methanol for 10 min, washed with PBS three times, incubated with 0.1% Triton X-100 in PBS for 15 min at room temperature, and washed again with PBS three times. Coverslips were blocked with 2% BSA in PBS for 1 hr before incubation with primary antibody (diluted 1:200 in 0.1% BSA in PBS) for 1 hr at room temperature. The coverslips were then washed three times with PBS before applying the secondary antibody (FITC-to-mouse IgG diluted 1:200 in 0.1% BSA in 1xPBS) for 1 hr. The coverslips were mounted with ProLong™ Glass Antifade Mountant with NucBlue™ Stain (Thermo Fisher, P36983). After 24 hours, imaging were performed with DAPI and GFP channels with an Olympus 100X objective (UPLSAPO, NA = 1.4).

### RT-qPCR

2 μg of the digested RNA was then reverse transcribed to cDNA in a 25 μL reaction using M-MLV Reverse Transcriptase (Promega M170A) with oligo dT following the manufacturer’s protocol. cDNA was diluted 10-fold with DEPC-treated water and RT-qPCR was performed using Luna Universal qPCR Master Mix (Biolabs, M3003E). RT-qPCR reactions were performed in 10 μL reactions that contained 4 μL of diluted cDNA, 5 μL 2x Luna Universal qPCR Master Mix, and 1 μL 5 μM gene-specific primer pairs. Primer sequences are as follows:

Timp2_D_F: TTTCTTGACATCGAGGACCC

Timp2_D_R: TCCAGGAAGGGATGTCAAAG

Timp2_P_F: ATGTGCGTGCTGGAATATGA

Timp2_P_R: CTGATACAGAGCATCAGGCG

Rbm39_F: GGTTCCAAGTAGACGATGAAGG

Rbm39_R: TGACAGTAGACCCTTGCCTCT

Pcf11_F: GGAAGAGAATATCTCACTGCCTT

Pcf11_R: GCAGAGGTTTAATAGGCCAAGC

GAPDH_F: TCACCACCATGGAGAAGGC

GAPDH_R: GCTAAGCAGTTGGTGGTGCA

Using the QuantStudio 5 Real-Time PCR System (Applied Biosystems) and 96-well white plates, the following cycling conditions were used: 95°C for 1 min, 40 amplification cycles of 95°C for 15 sec followed by 60°C for 30 sec, and a final melting cycle of 95°C for 1 ses, 60°C for 20 sec, and 95°C for 1 s. Subsequently, a melt curve was performed to verify that amplified products were a single discrete species. Threshold cycle (CT) values were automatically calculated by the LightCycler system and relative transcript levels (compared to mouse GAPDH) were calculated using the 2^−ΔΔCT^ method. RT-qPCR reactions were performed using three independent biological replicates, with each replicate having two technical replicates.

### Cell proliferation assay

NIH3T3 cells were seeded in 24-well plates at 1 x 10^4^ cells per well. For each cell line, at least four wells were used for each experiment. Cells were counted and cell number per well was calculated after 24 hrs and 48 hrs post seeding (N1 and N2). The cell doubling time (Td) can be calculated by:

### Cell migration assay

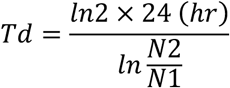

NIH3T3 cells were grown in a 6-well plate or 12-well plate to full confluence. A P200 pipette tip was used to make a scratch in each well. Cell debris were removed by washing with 1 x PBS twice. Cells were then cultured in DMEM + 0.5% (v/v) calf serum. A total of 3-5 points along each scratch were marked and photos were taken at each of the marked positions to record the original boundary of scratch in the EVOS FL microscopic system. Photos were then taken after 18-24 hrs to record cell migration.

### Focus formation assay

For each cell line, 2 x 10^4^ cells were seeded in a 6-cm dish and cultured for 10 days. Media was changed every 2 days. At day 10, cells were washed twice with 1 x PBS and fixed with ice-cold methanol on an ice box for 10 min. The cells were then stained with 0.5% (w/v) crystal violet solution (25% methanol) for 10 sec and rinsed with water multiple times until there was no dye coming out of the dishes. Dished were air-dried at room temperature overnight before imaging.

### Single-molecule FISH

#### Design and generation of smFISH probes

smFISH probes were designed by and ordered from Stellaris (now part of Biosearch Technologies). The full list of probe sequences is in the **Supplementary Table S6**.

#### Hybridization

NIH3T3 cells were seeded and cultured on coverslips in 12-well plates for 24 hrs. To fix samples, cells were washed twice with PBS, fixed with Fixation buffer (3.7% (w/v) formaldehyde in 1xPBS) for 15 min, and kept at 4 °C in 70% ethanol. Before hybridization, 70% ethanol was removed from wells and cells were washed with Wash buffer (3.7% (w/v) formaldehyde, 0.3 M NaCl, 0.03 M sodium citrate) for at least 2 min. For each coverslip, 100 µL of Hybridization buffer (10% (w/v) dextran sulfate, 3.7% (w/v) formaldehyde, 0.3 M NaCl, 0.03 M sodium citrate) with 2 µL 25 ug/uL smFISH probes was used. Coverslips were put on a piece of parafilm into a sealed cell culture dishes to avoid evaporation. After overnight hybridization in a 37 °C dark incubator, coverslips were washed twice, 30 min each, in the Wash buffer. For the second wash, DAPI was added directly into the Wash buffer for staining of the cell nucleus. The coverslip was then washed with 2xSSC for three times and then submerged in Antifade buffer (0.3 M NaCl, 0.03 M sodium citrate, 0.4 (w/v) glucose, 0.01 M Tris). For mounting of coverslips, 100 µL antifade buffer with addition of 1 µL Glucose oxidase (Catalog # G2133-10KU, Sigma) and 1 µL Catalase (Catalog # C3515-10MG, Sigma) was used. The edge of each coverslip was sealed with nail polish.

#### Imaging

smFISH samples were imaged on an Olympus IX83 microscope. Z stack intervals were set to 0.3 µm. Images were captured by using an Andor iXon Life 888 EMCCD camera with 1,024 × 1,024 pixels and a dynamic range based on 16-bit digits. An Olympus 100X objective (UPLSAPO, NA = 1.4) was used for imaging. Fluorescent signals were detected by using the following filters: Excitation = 599/13 nm and Emission = 632/28 nm for probes labeled with TexasRed, Excitation = 640/30 nm and Emission = 690/50 nm for probes labeled with Q670. The Olympus CellSens software was used for image acquisition and initial data processing. Images in .tiff format in separate channels were exported for further analysis.

#### smFISH data analysis

We used rajlabformattools (https://github.com/arjunrajlaboratory/rajlabformattools) and rajlabimagetools (https://github.com/arjunrajlaboratory/rajlabimagetools) following the instructions on the website for analysis our smFISH data ^38^. Briefly, all channels of images were stacked together for manual cell segmentation. Within the area range of each cell, signal dots were identified by an algorithm using Gaussian fit over the z-stack images within each channel, and the exact number of dots reflecting the number of mRNA molecules was counted. Then we performed co-localization analysis for the TexasRed and Q670 channels to identify the number of *Timp2* long mRNA (dots that were colocalized between the two channels) and short mRNA. For the two groups of dots, we calculated their center position and their average distance to the center, and the ratio of the two average distances were defined as Relative Distribution Index (RDI).

### Luciferase assays

NIH3T3 cells were seeded at 1 x 10^5^ cells per well in 24-well plates. For each well, 25 ng of pGS or pRF vector was transfected into cells by using Lipofectamine 3000 (Invitrogen, 2292027) according to the manufacturer’s instructions. Media was changed 6 hrs post transfection.

#### Gaussia / SEAP luciferase assay

For samples transfected with the pGS vector for Gaussia and SEAP luciferase assays, media was collected 18-24 hrs post transfection. Pierce™ Gaussia Luciferase Glow Assay Kit (Catalog # 16160, Thermo Fisher scientific) and Phospha-Light™ SEAP Reporter Gene Assay Kit (Catalog # T1015, Thermo Fisher scientific) were used for measurements of Gaussia luciferase and SEAP activities.

#### Renilla / Firefly luciferase assay

For samples transfected with the pRF vector for Renilla and firefly luciferase assays, cell lysates were obtained 24 hrs post transfection and were used for measurements of Renilla and firefly luciferases by using the Luc-Pair Duo-Luciferase Assay Kit 2.0 (Catalog # LF001, GeneCopoeia,) according to manufacturer’s instruction.

### Quantification and statistical analysis

Student t-test was used for comparison of paired samples. For all panels with more than two samples, statistical significance for difference was assessed by one-way ANOVA. Statistical details and error bars are defined in the legend of each figure.

## ACKNOWLEDGEMENTS

We thank Wei Lu for her contributions at the early stage of this work. We thank Margaret Dunagin and Arjun Raj for technical assistance of smFISH. Y.A. was supported by the American Heart Association (827222) and the Blavatnik Family Foundation. A.I. was supported by NIH grant F31 CA277953. M.M. was supported by NIH grant R01 CA102184. B.T. was supported by NIH grants (R01GM084089 and R35GM153277).

## AUTHOR CONTRIBUTIONS

Y.A. and B.T. conceived and designed the project. Y.A., Q.D. and Z.W. performed experiments and analyzed data. S.T., A.I., and M.M. provided guidance on experiments. Y.A. and B.T. wrote the manuscript with input from all authors.

