## Supplementary material for "Regulation of alternative polyadenylation isoforms of *Timp2* is an effector event of RAS signaling in cell transformation"

Figure S1

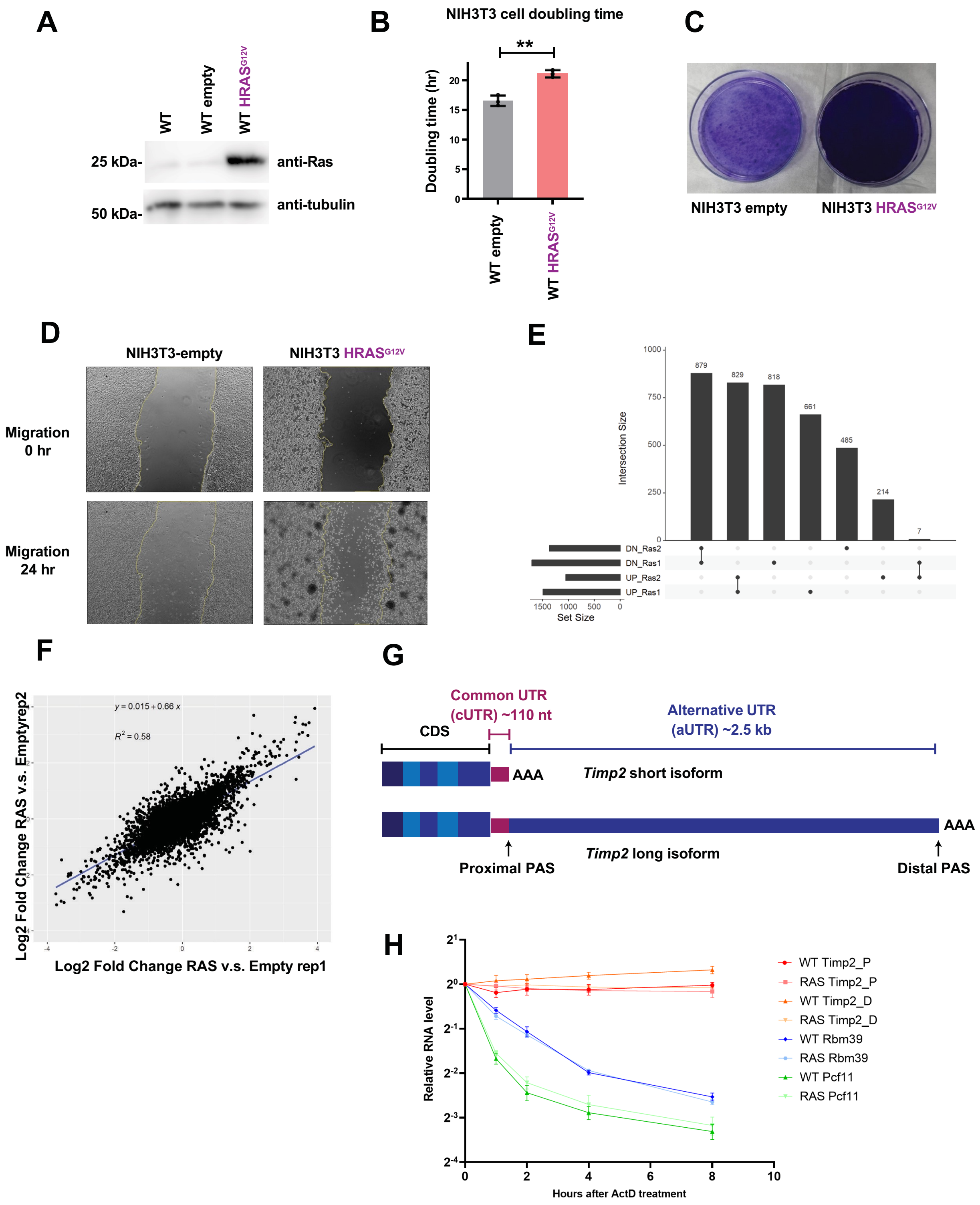

**Supplementary Figure S1. (A)** Western blotting result showing RAS protein level in NIH3T3 WT, NIH3T3 -empty and NIH3T3-HRAS$^{G12V}$ cells. **(B)** Cell doubling time analysis of NIH3T3 -empty and NIH3T3-HRAS$^{G12V}$ cells. **(C)** Focus formation assay of NIH3T3 -empty and NIH3T3-HRAS$^{G12V}$ cells. **(D)** Scratch-based cell migration assay for NIH3T3 -empty and NIH3T3-HRAS$^{G12V}$ cells over a 24 hr period. **(E)** UpSet plot showing number of commonly regulated genes in two QuantSeq-Pool replicates for NIH3T3-HRAS$^{G12V}$ cells vs. NIH3T3-empty cells. **(F)** Linear regression fit and $R^2$ for the scatter plot in **Figure 1B**. **(G)** Diagram of *Timp2* short and long 3'UTR isoforms. **(H)** RNA stability assay. Actinomycin D was added to WT and HRAS$^{G12V}$ cells for 0, 1, 2, 4 and 8 hours before total RNA was collected. RT-qPCR were used to measure the relative RNA level to Gapdh.

Figure S2

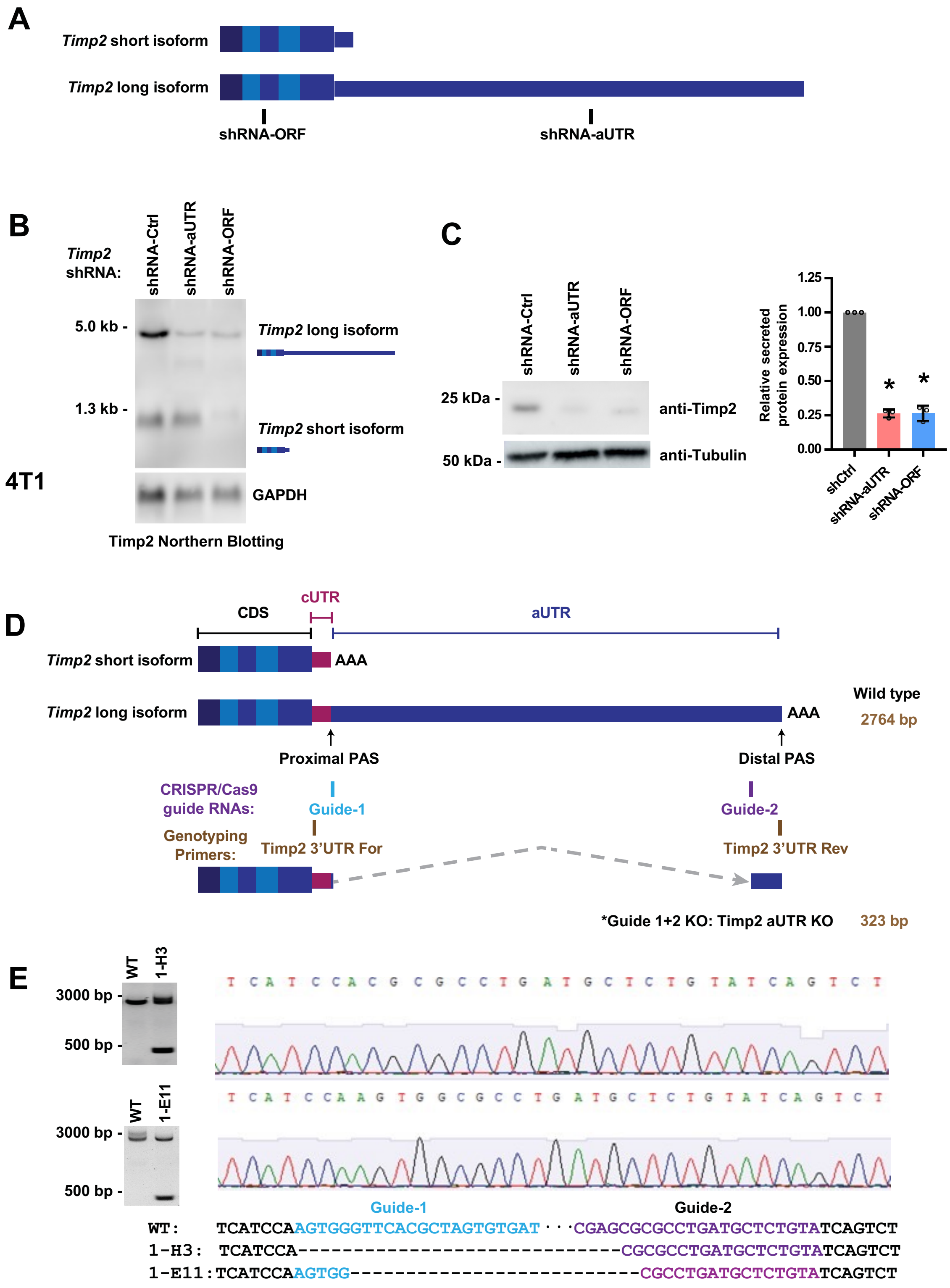


**Supplementary Figure S2. (A)** Diagram showing the shRNA targeting site on *Timp2* transcripts used in **Figure 2**. **(B)** Northern blotting result showing *Timp2* mRNA level in 4T1 cells expressing shRNA-Ctrl, shRNA-aUTR (knockdown of only the Timp2 long mRNA isoform), or shRNA-ORF (knockdown of both *Timp2* short and long mRNA isoforms). **(C)** Western blotting result showing secreted Timp2 protein level of 4T1 cells expressing shRNA-Ctrl, shRNA-aUTR or shRNA-ORF. Data quantifications are also presented. **(D)** Diagram showing CRISPR/Cas9 guide RNA targeting sites in the genomic sequence of *Timp2* 3'UTR. Guide-1 is in blue and guide-2 in purple. Genotyping primers are in brown. The WT sequence should give rise to a PCR product of 2,764 bp by using the genotyping primers, whereas the aUTR KO sequence should give rise to a PCR product of 323 bp by using the same pair of primers. **(E)** Genotyping and Sanger sequence results of 1-H3 and 1-E11 cell lines.

Figure S3

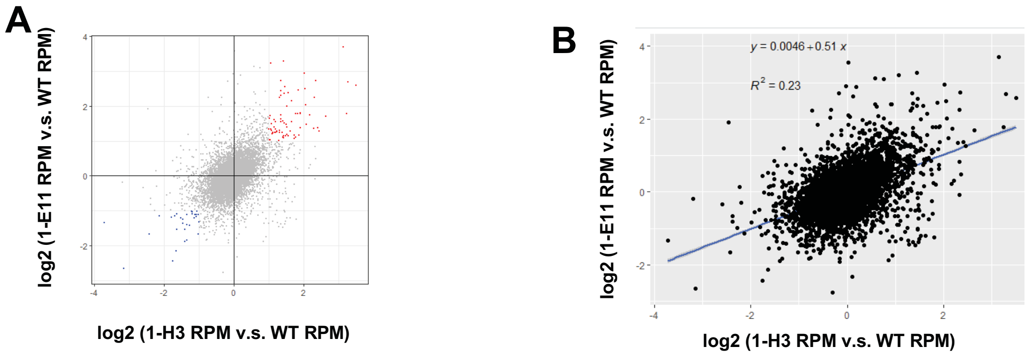


Figure S3 (continued)

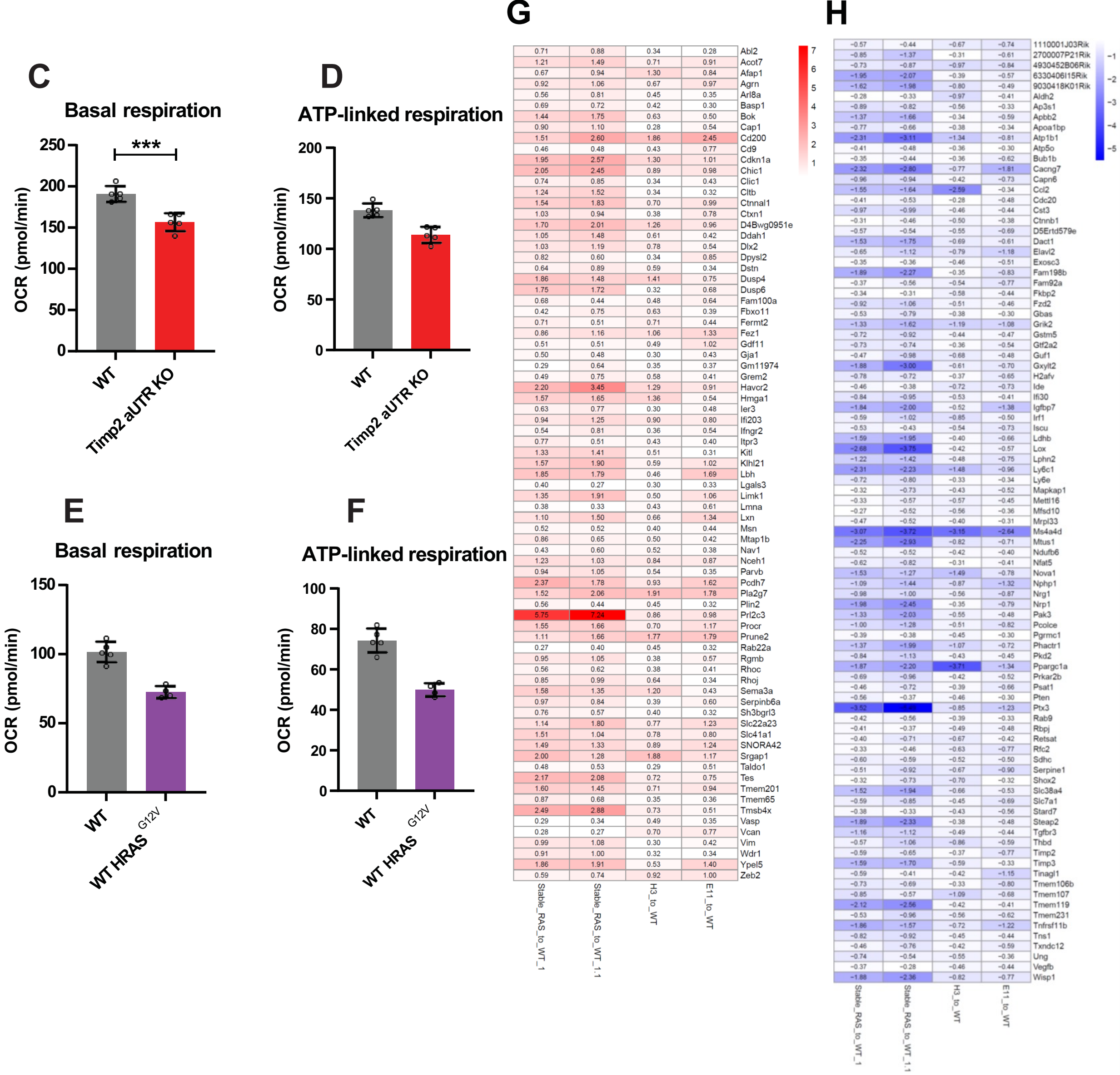


**Supplementary Figure S3. (A)** Scatter plot for genes shown in **Figure 3A** and **3B**. **(B)** Linear regression fit and $R^2$ for the scatter plot in (A). **(C)** Basal respiration rates of NIH3T3 *Timp2*

aUTR KO$^{+/-}$ cells and WT cells as measured by the Seahorse assay. **(D)** ATP-linked respiration rates of NIH3T3 *Timp2* aUTR KO$^{+/-}$ cells and WT cells as measured by the Seahorse assay. **(E)** Basal respiration rates of NIH3T3-HRAS$^{G12V}$ cells and NIH3T3-empty cells as measured by the Seahorse assay. **(F)** ATP-linked respiration rates of NIH3T3-HRAS$^{G12V}$ cells and NIH3T3-empty cells as measured by the Seahorse assay. **(G)** The 77 commonly upregulated genes shown in **Figure 3F**. (H) The 91 commonly downregulated genes shown in **Figure 3F**.

Figure S4

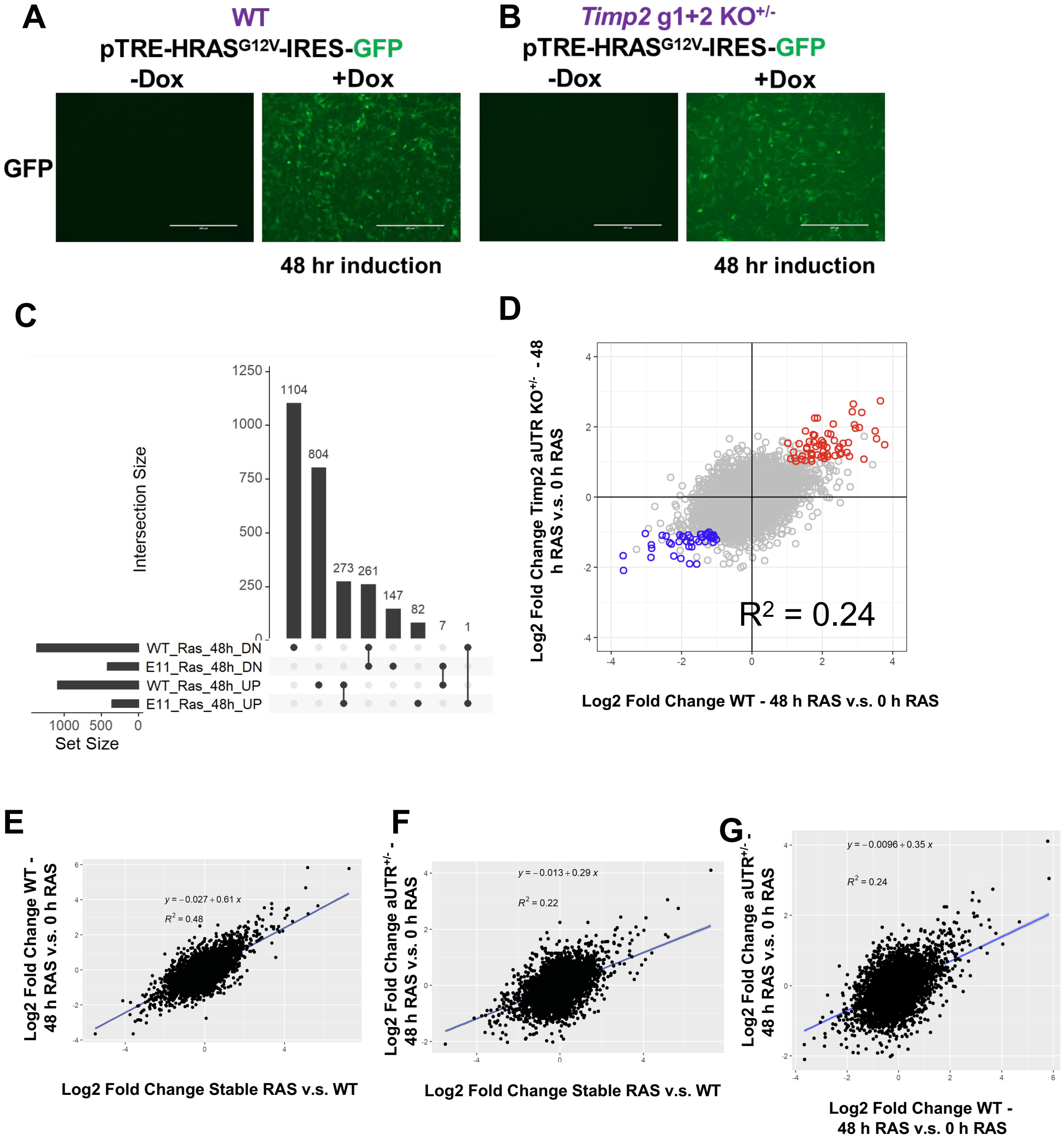

A
WT
pTRE-HRASG12V-IRES-GFP
-Dox
+Dox
GFP
48 hr induction
B
Timp2 g1+2 KO+/-
pTRE-HRASG12V-IRES-GFP
-Dox
+Dox
48 hr induction
C
Intersection Size
1104
804
273
261
147
82
7
1
WT_Ras_48h_DN
E11_Ras_48h_DN
WT_Ras_48h_UP
E11_Ras_48h_UP
Set Size
D
Log2 Fold Change Timp2 aUTR KO+/- - 48 h RAS v.s. 0 h RAS
R2 = 0.24
Log2 Fold Change WT - 48 h RAS v.s. 0 h RAS
E
Log2 Fold Change WT - 48 h RAS v.s. 0 h RAS
y = −0.027 + 0.61 x
R2 = 0.48
Log2 Fold Change Stable RAS v.s. WT
F
Log2 Fold Change aUTR+/- - 48 h RAS v.s. 0 h RAS
y = −0.013 + 0.29 x
R2 = 0.22
Log2 Fold Change Stable RAS v.s. WT
G
Log2 Fold Change aUTR+/- - 48 h RAS v.s. 0 h RAS
y = −0.0096 + 0.35 x
R2 = 0.24
Log2 Fold Change WT - 48 h RAS v.s. 0 h RAS

Figure S4
(continued)

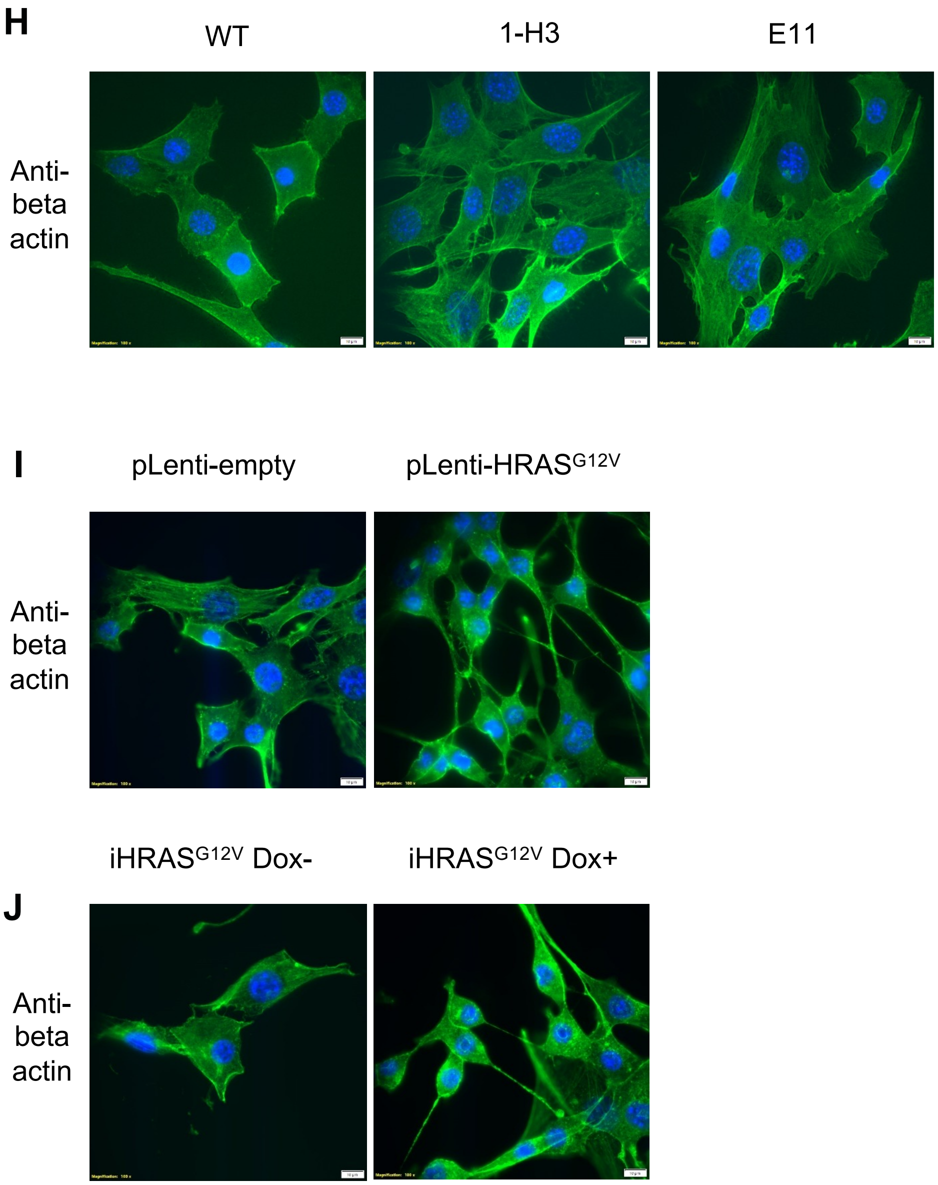


**Supplementary Figure S4. (A)** Dox induction of expression of HRAS$^{G12V}$-IRES-GFP in NIH3T3 WT cells. **(B)** Dox induction of expression of HRAS$^{G12V}$-IRES-GFP in NIH3T3 aUTR KO$^{+/-}$ cells.

**(C)** UpSet plot showing commonly regulated genes between (1) inducible $HRAS^{G12V}$ vs. WT cells and (2) inducible $HRAS^{G12V}$ vs. aUTR $KO^{+/-}$ cells. **(D)** Scatter plot of commonly regulated genes identified in (C). **(E)** Linear regression fit and $R^2$ for the scatter plot in **Figure 4C**. **(F)** Linear regression fit and $R^2$ for the scatter plot in Figure 4E. **(G)** Linear regression fit and $R^2$ for the scatter plot in **Figure S4D**. **(H)** Immunofluorescence of beta-actin in WT and *Timp2* aUTR $KO^{+/-}$ cells. **(I)** Immunofluorescence of beta-actin in WT-empty and $HRAS^{G12V}$ cells. **(J)** Immunofluorescence of beta-actin in $iHRAS^{G12V}$ cells.

Figure S5

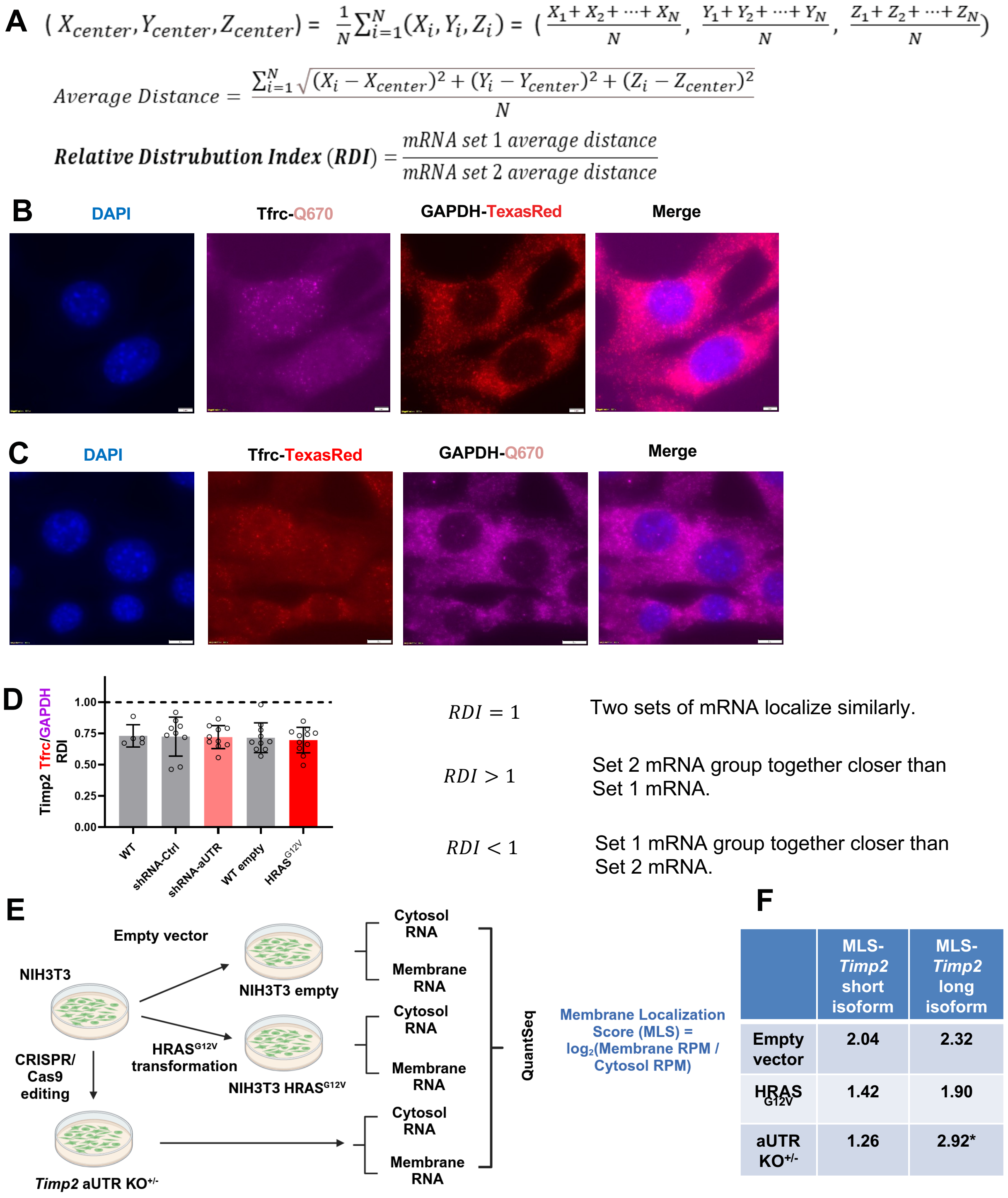


| | MLS-*Timp2* short isoform | MLS-*Timp2* long isoform |
|---|---|---|
| Empty vector | 2.04 | 2.32 |
| HRAS G12V | 1.42 | 1.90 |
| aUTR KO+/- | 1.26 | 2.92* |

**Supplementary Figure S5. smFISH data for *Tfrc* and *Gapdh* and the cell fractionation assay. (A)** Calculation details for relative distribution index (RDI). **(B)** smFISH for *Tfrc* and

*Gapdh* with *Tfrc* labeled with Quasar 670 (Q670) and *Gapdh* with TexasRed. Pseudo colors are shown. **(C)** smFISH for *Tfrc* and *Gapdh* with *Tfrc* probes labeled with TexasRed and *Gapdh* probes with Quasar670. Pseudo colors are shown. **(D)** RDI calculated for *Tfrc* and *Gapdh* across five NIH3T3 cell lines used for further smFISH studies in **Figure 5**. **(E)** Diagram showing samples and procedures of cell fractionation and the QuantSeq-Pool library preparation for those samples. **(F)** Membrane localization score (MLS) calculated for samples shown in **E**.

Figure S6

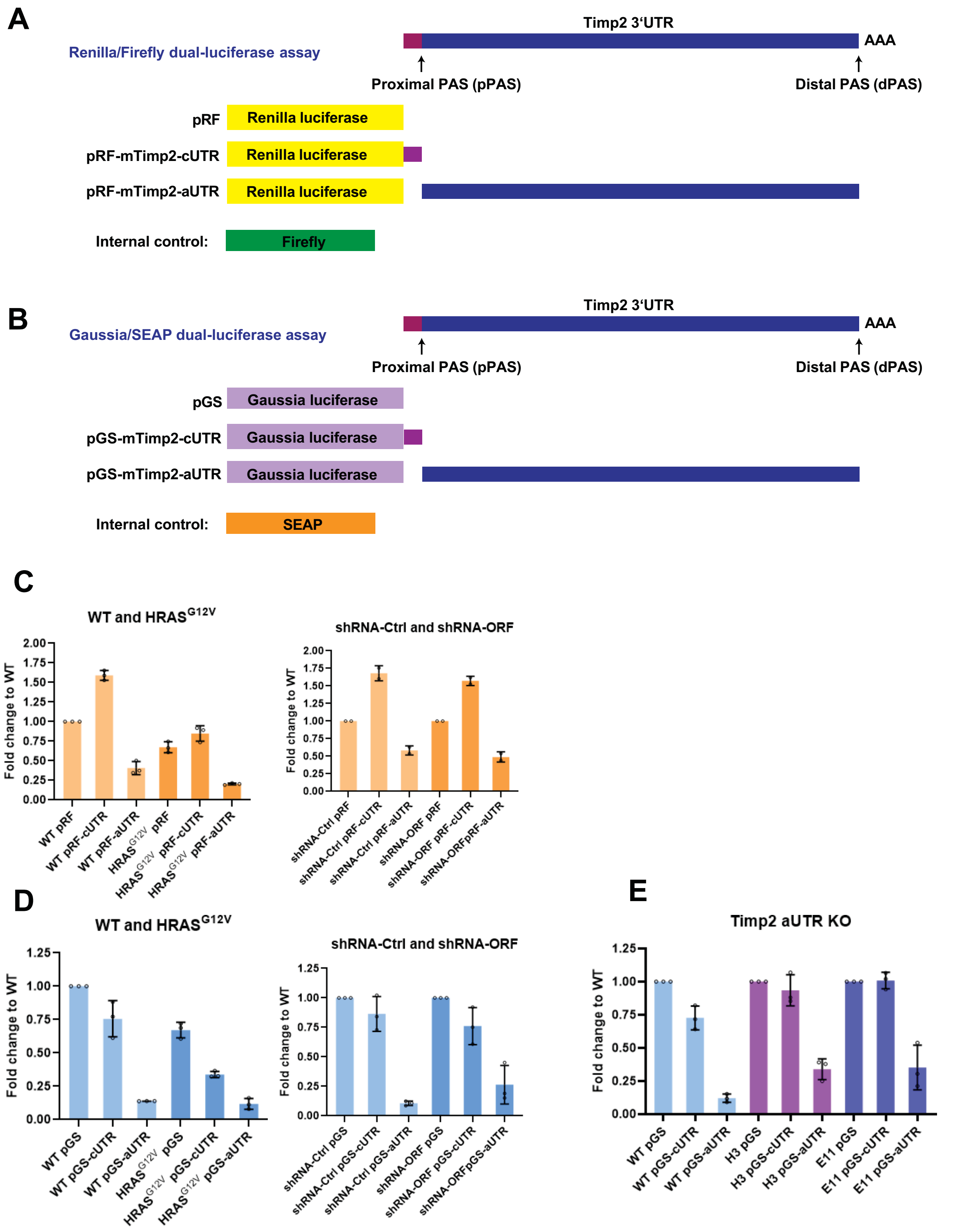


**Supplementary Figure S6. (A)** Diagram of the set of vectors used in the Renilla and firefly luciferase assays. **(B)** Diagram of the set of vectors used in the Gaussia luciferase and SEAP assays. **(C)** Raw data of the Renilla and firefly luciferase assays for each vector in NIH3T3-empty and NIH3T3-HRAS$^{G12V}$ cells (left) as well as NIH3T3 cells expressing shRNA-Ctrl or shRNA-ORF (right). **(D)** Raw data of Gaussia luciferase and SEAP assay for each vector in NIH3T3-empty and NIH3T3-HRAS$^{G12V}$ cells (left) as well as NIH3T3 cells expressing shRNA-Ctrl or shRNA-ORF cells (right). **(E)** Raw data of the Gaussia luciferase and SEAP assays for each vector in NIH3T3 WT, 1-H3 and 1-E11 cells.

**Supplementary Table S1: smFISH probe sequence**

All probes are modified with 3'-end mdC(TEG-Amino).

| | name | oligo sequence |
|---|---|---|
| probes for mouse Tfrc | mTfrc_1 | aatgctgatctggcttgatc |
| | mTfrc_2 | aaggctaaaccgggtgtatg |
| | mTfrc_3 | tcatctccacatgactgtta |
| | mTfrc_4 | tagccttcatgttattgtcg |
| | mTfrc_5 | aaacctcttgggttttctga |
| | mTfrc_6 | gtgcaatagctgcaaagcag |
| | mTfrc_7 | tctacacgcttacaatagcc |
| | mTfrc_8 | ttgaggtctgcccaatataa |
| | mTfrc_9 | ggagttcaacttctctgaca |
| | mTfrc_10 | aggagtgtatgtattctggc |
| | mTfrc_11 | tcatctttttgagatccagc |
| | mTfrc_12 | gccagactttgctgaattta |
| | mTfrc_13 | ctatggtcaccatgttttga |
| | mTfrc_14 | actgggtctaagttaccatt |
| | mTfrc_15 | gtaggtttactgaatgccac |
| | mTfrc_16 | ggaccagtttaccagaaact |
| | mTfrc_17 | ctgcaaaagtaatttcccct |
| | mTfrc_18 | ttgcattaaagctttgggca |
| | mTfrc_19 | tccaaagagtgcaaggtctg |
| | mTfrc_20 | tattaggcaaccctgatgac |
| | mTfrc_21 | atctatgttccatctagcag |
| | mTfrc_22 | ttctgtgaaagttccagctt |
| | mTfrc_23 | gggctcctactacaacataa |
| | mTfrc_24 | ttcaacagaagacctgttcc |
| | mTfrc_25 | aagatgaaaggtatccctcc |
| | mTfrc_26 | ctaatccaattgctgtctct |
| | mTfrc_27 | ctgggattccagaatatgca |
| | mTfrc_28 | tctgagtcaatgcctcatag |
| | mTfrc_29 | aaccatttggttgagctgag |

| | | |
|---|---|---|
| | mTfrc_30 | atccctgatatctgttttga |
| | mTfrc_31 | ggaatacagccactgtagac |
| | mTfrc_32 | agtagcacggaagtagtctc |
| | mTfrc_33 | atagggcgacaggaagtgat |
| | mTfrc_34 | tgtcggaaaggagactctct |
| | mTfrc_35 | gttctccactaaagctgaga |
| | mTfrc_36 | tccaaatgtcaccagagagg |
| probes for mouse Gapdh | mGapdh_1 | caaatccgttcacaccgac |
| | mGapdh_2 | aaatggcagccctggtgac |
| | mGapdh_3 | caatctccactttgccact |
| | mGapdh_4 | tgaaggggtcgttgatggc |
| | mGapdh_5 | agaccatgtagttgaggtc |
| | mGapdh_6 | ccgtgagtggagtcatact |
| | mGapdh_7 | cttgactgtgccgttgaat |
| | mGapdh_8 | gatgacaagcttcccattc |
| | mGapdh_9 | ggaagatggtgatgggctt |
| | mGapdh_10 | accccatttgatgttagtg |
| | mGapdh_11 | tccacgacatactcagcac |
| | mGapdh_12 | tctccatggtggtgaagac |
| | mGapdh_13 | cggagatgatgacccttt |
| | mGapdh_14 | acaaacatgggggcatcgg |
| | mGapdh_15 | atttctcgtggttcacacc |
| | mGapdh_16 | gcattgctgacaatcttga |
| | mGapdh_17 | taagcagttggtggtgcag |
| | mGapdh_18 | ttgtcatggatgaccttgg |
| | mGapdh_19 | catgagcccttccacaatg |
| | mGapdh_20 | agtgatggcatggactgtg |
| | mGapdh_21 | atccacagtcttctgggtg |
| | mGapdh_22 | gggatgatgttctgggcag |
| | mGapdh_23 | ttggcagcaccagtggatg |
| | mGapdh_24 | cgttcagctctgggatgac |
| | mGapdh_25 | ggggtaggaacacggaagg |
| | mGapdh_26 | agatccacgacggacacat |

| | | |
|---|---|---|
| | mGapdh_27 | catacttggcaggtttctc |
| | mGapdh_28 | cttcaccaccttcttgatg |
| | mGapdh_29 | caagatgcccttcagtggg |
| | mGapdh_30 | aacctggtcctcagtgtag |
| | mGapdh_31 | gagttgctgttgaagtcgc |
| | mGapdh_32 | cggcatcgaaggtggaaga |
| | mGapdh_33 | gtcattgagagcaatgcca |
| | mGapdh_34 | ccaggaaatgagcttgaca |
| | mGapdh_35 | gtagccgtattcattgtca |
| | mGapdh_36 | atgtaggccatgaggtcca |
| probes for mouse Timp2 coding region and the common 3'UTR | Timp2-P_35 | tccgaggatcctctgctc |
| | Timp2-P_56 | cgagccgccgttttattg |
| | Timp2-P_84 | tatggctgtgtttgctgc |
| | Timp2-P_156 | cacccggcggagaagaaa |
| | Timp2-P_200 | ctttgtcaaggggcgcag |
| | Timp2-P_219 | cgcgcaaactttctgtcc |
| | Timp2-P_361 | cgtggctagcagcaggag |
| | Timp2-P_419 | aaaacgcctgttgcgggt |
| | Timp2-P_437 | tcactacgtctgcattgc |
| | Timp2-P_455 | tcactgctttggctctga |
| | Timp2-P_474 | gaatccacctccttctcg |
| | Timp2-P_492 | ccatagatgtcattcccg |
| | Timp2-P_510 | atcctcttgatggggttg |
| | Timp2-P_530 | tctgcttgatctcatact |
| | Timp2-P_560 | tgtctttgtcaggtcctt |
| | Timp2-P_582 | ggggccgtgtagataaac |
| | Timp2-P_600 | ccgcacactgctgaagag |
| | Timp2-P_619 | tccaacgtccagcgagac |
| | Timp2-P_637 | tagatactccttctttcc |
| | Timp2-P_655 | ttctgcctttcctgcaat |
| | Timp2-P_675 | atgtgcatcttgccatct |
| | Timp2-P_700 | gggcacaatgaagtcaca |
| | Timp2-P_718 | gatgctaagcgtgtccca |

| | | |
|---|---|---|
| | Timp2-P_736 | caggctcttcttctgggt |
| | Timp2-P_754 | catctggtacctgtggtt |
| | Timp2-P_772 | gatcttgcactcacagcc |
| | Timp2-P_792 | gggatcatgggacagcga |
| | Timp2-P_810 | ggggaggagatgtagcaa |
| | Timp2-P_828 | atccagaggcactcatcc |
| | Timp2-P_846 | ttctctgtgacccagtcc |
| | Timp2-P_864 | tggtgcccattgatgctc |
| | Timp2-P_882 | caggcgaagaacttggcc |
| | Timp2-P_900 | ccatcacttctcttgatg |
| | Timp2-P_918 | cggtaccacgcgcaagaa |
| | Timp2-P_945 | aactcttgcttggggggt |
| | Timp2-P_963 | gggtcctcgatgtcaaga |
| | Timp2-P_981 | ctctgtcagccttcttac |
| | Timp2-P_999 | ttcaattggccacagggg |
| | Timp2-P_1019 | gtctaaaccctcagaggc |
| probes for mouse Timp2 alternative 3'UTR | Timp2-D_87 | tgcctgtagcaaggatcaaa |
| | Timp2-D_117 | agacctggtaacaagtctgt |
| | Timp2-D_203 | tgcaaaatgcacattcccag |
| | Timp2-D_239 | cggctacacagtcttacaac |
| | Timp2-D_275 | acatgtcactctcaggatga |
| | Timp2-D_330 | atgggaagcatttgcaaggg |
| | Timp2-D_356 | gaaagtccataccagatgca |
| | Timp2-D_397 | gttggagggtgattcttaga |
| | Timp2-D_429 | ctgactgggactcctagaaa |
| | Timp2-D_450 | gaagtctgtggattcatggg |
| | Timp2-D_493 | ggttgaagccagaaacacgg |
| | Timp2-D_513 | gaggaaggcagcaatgactt |
| | Timp2-D_537 | gagggtgtgtgtagagacag |
| | Timp2-D_606 | caacaaggactgccaagcac |
| | Timp2-D_631 | aatggcttgggaagcttgag |
| | Timp2-D_652 | cttgcttggaaaggggtgaa |
| | Timp2-D_705 | tgatgcgacagagagccaaa |

| | | |
|---|---|---|
| | Timp2-D_753 | tgtgcaaaagagggagtgct |
| | Timp2-D_774 | tggccttttatatcatcagt |
| | Timp2-D_816 | tagcagggagagttcactag |
| | Timp2-D_841 | aacagggtcataatgaccgc |
| | Timp2-D_867 | cgtgcttagttactgacaca |
| | Timp2-D_986 | tcagccatgcatgagaattc |
| | Timp2-D_1042 | tataagggccttaagcgaca |
| | Timp2-D_1062 | aagtagatataccagttccc |
| | Timp2-D_1133 | gttagtgataaagctctgcc |
| | Timp2-D_1203 | tatacatttgtgtccttggg |
| | Timp2-D_1311 | gttggagcaaagaggctttt |
| | Timp2-D_1337 | ttggggtttctgatggttag |
| | Timp2-D_1411 | ttaaaaggactggccaggtc |
| | Timp2-D_1474 | tgcacaacacgaaaatgccc |
| | Timp2-D_1498 | ccagcatccaaggaagaaca |
| | Timp2-D_1555 | acaggaagcaaacaggcagg |
| | Timp2-D_1593 | cagcagaaacaggcactgtt |
| | Timp2-D_1643 | aagaacgctcacggcgaaca |
| | Timp2-D_1663 | gaaaggcatcaacacggcac |
| | Timp2-D_1792 | aagatgagaaagggtccctc |
| | Timp2-D_1831 | ggagaggctggaaaagcaga |
| | Timp2-D_1884 | gtatactcagagggtctttc |
| | Timp2-D_1906 | gaaaggccgtctttcagaac |
| | Timp2-D_1926 | cccagcatgagtggaaaaca |
| | Timp2-D_2044 | caattggtttgctgggaagg |
| | Timp2-D_2139 | aaatattgcgagttgctggc |
| | Timp2-D_2175 | gcgatttggaaattgctacc |
| | Timp2-D_2196 | tgtcttaatacgcatcaggc |
| | Timp2-D_2310 | cagacttcatattccagcac |
| | Timp2-D_2371 | ggcttgctgactttcaacaa |
| | Timp2-D_2397 | actgatacagagcatcaggc |
